# Septin Roles and Mechanisms in Organization of Endothelial Cell Junctions

**DOI:** 10.1101/2020.03.04.977199

**Authors:** Joanna Kim, John A. Cooper

**Affiliations:** Department of Biochemistry & Molecular Biophysics, Washington University School of Medicine, Saint Louis, MO, USA

## Abstract

Septins play an important role in regulating the barrier function of the endothelial monolayer of the microvasculature. Depletion of septin 2 protein alters the organization of vascular endothelial (VE)-cadherin at cell-cell adherens junctions as well as the dynamics of membrane protrusions at endothelial cell-cell contact sites. Here, we report the discovery that localization of septin 2 at endothelial cell junctions is important for the distribution of a number of other junctional molecules. We also found that treatment of microvascular endothelial cells with the inflammatory mediator TNF-α led to sequestration of septin 2 away from cell junctions and into the cytoplasm, without an effect on the overall level of septin 2 protein. Interestingly, TNF-α treatment of endothelial monolayers produced effects similar to those of depletion of septin 2 on various molecular components of adherens junctions (AJs) and tight junctions (TJs). Immunofluorescence staining revealed disruption of the integrity of AJs and TJs at cell-cell junctions without significant changes in protein expression except for VE-cadherin and nectin-2. To investigate the mechanism of junctional localization of septin 2, we mutated the polybasic motif of septin 2, which is proposed to interact with PIP_2_ in the plasma membrane. Overexpression of PIP_2_-binding mutant (PIP_2_BM) septin 2 led to loss of septin 2 from cell junctions with accumulation in the cytoplasm. This redistribution of septin 2 away from the membrane led to effects on cell junction molecules similar to those observed for depletion of septin 2. We conclude that septin localization to the membrane is essential for function and that septins support the localization of multiple cell junction molecules in endothelial cells.

## Introduction

The septin family of genes and proteins were discovered as cytoskeletal elements important for cell division in budding yeast ^1^. Septins form filaments that assemble into a ring associated with the cell membrane at the site of cytokinesis ^2,3^. Septins are found in a wide variety of eukaryotes and cell types, where they play a diverse set of roles in cellular processes ^4–6^. Septin family proteins are well-conserved across species, and mammalian septins consist of 13 members classified into 4 different groups on the basis of sequence similarity and domain structure ^7–9^.

Septin family proteins form homo-and hetero-oligomers *in vitro* and *in vivo* ^10,11^. The oligomers assemble into filaments and associate with membranes via a polybasic domain ^12–14^, and the filaments assemble further into ring- and gauze-shaped higher-order arrays *in vitro* and in cells ^3,15,16^. Thus, septin filaments represent an important element of the membrane cytoskeleton, and they contribute to membrane-associated cellular functions ^10,11,17^. For instance, septin filaments are enriched at positively curved areas of plasma membrane, providing interactive feedback between cell shape and the cytoskeleton ^10,18–20^. In addition, septins play roles in intracellular membrane-associated processes ^21–23^, such as mitochondrial fission ^24^ and endomembrane fusion ^25^. Septin filaments interact with other cytoskeletal elements, including actin filaments and microtubules, providing crosstalk regulation of cell morphology, migration, and other functions in cooperative manners in multiple cell settings ^26–31^.

The endothelial monolayer is a continuous thin layer of endothelial cells that lines the interior surface of blood and lymphatic vessels. The endothelial monolayer is an active and regulated barrier ^32^, playing crucial roles in multiple biological processes, including vascular tone, thrombosis / thrombolysis, cell adhesion and passage of small molecules and cells. In order to perform these diverse roles, maintenance and regulation of the barrier structure and monolayer integrity are crucial ^33–35^. Endothelial cell-cell junctions are key components in regulating the integrity of the endothelial monolayer, and they consist of adherens junctions, tight junctions, and gap junctions, along with other diffusely localized cell adhesion proteins ^36–38^.

Adherens junctions contain vascular endothelial cadherin (VE-cadherin), nectin, and afadin. VE-cadherin, the dominant component of adherens junctions, is a transmembrane glycoprotein that forms homodimers via its extracellular domain. The cytosolic domain of VE-cadherin interacts with catenin complexes (p120, β-catenin, and α-catenin) linking actin filaments to the junctions ^39^. Afadin interacts with actin filaments and nectin as part of its recruitment to VE-cadherin-based adherens junctions ^40^. Afadin is also recruited to tight junctions ^40,41^, the closely apposed junctional areas assembled by intercellular adhesion molecules including claudins, occludin, and zonula occludens (ZOs) ^42,43^. ZO proteins function as cytosolic scaffolds that directly bind to actin filaments and are recruited to lateral membranes, where they bind claudins and occludin to promote tight junction assembly ^44^. Like afadin, ZO proteins have shared functions to promote the assembly of both tight junction and adherens junction ^45^. Junctional adhesion molecules (JAMs) are another set of intercellular adhesion components that are located at tight junctions and regulate interactions between endothelial cells and between leukocytes and endothelial cells ^46^ . Gap junctions are comprised of members of the connexin protein family. Connexon (hexameric connexin complex) form channel-like structure between adjacent cells and regulate transferring intracellular molecules ^47,48^. In addition to these molecules, platelet endothelial cell adhesion molecule (PECAM-1, also known as CD31) ^49^ and MIC-2 (also known as CD99) are recruited to endothelial cell junctions, where they regulate transmigration of leukocytes across the junctions ^50,51^.

Cell junctions are supported in the cytoplasm by elements of the cytoskeleton associated with the plasma membrane, notably actin filaments (F-actin) ^52–56^. In previous work, we characterized septin filaments as a membrane cytoskeleton component of endothelial cells. Septins were found at positively curved membrane areas, near the base of actin-based lamellipodial protrusions. Loss of septins impaired the assembly and function of VE-cadherin-based adherens junctions and the overall integrity of endothelial monolayers ^20^. Here, we investigated whether and how septins influence the integrity of other cell junctional molecules, using the same endothelial cell monolayer system. We also investigated the mechanism of septin association with the membrane and the relationship of septin function to effects of the inflammatory mediator TNF-α on endothelial junctional integrity.

## Materials and Methods

### Cells and Cell Culture

Primary human dermal microvascular endothelial cells (HDMVECs) were cultured as described ^20^. In brief, primary human dermal microvascular endothelial cells (HDMVECs) from neonates, obtained from Lonza Bioscience (Walkersville, MD, USA), were grown in EGM-2MV (Microvascular Endothelial Cell Growth Medium-2) medium from Lonza Bioscience. HDMVECs were used between passages 3 and 8 for experiments. For fluorescence imaging, 5×10^4^ HDMVECs were seeded into 14-mm glass-bottom dishes (MatTek Corporation, Ashland, MA, USA) and cultured for two to three days until cells formed continuous monolayers. For infection of cells with lentivirus, 2×10^5^ cells were seeded into 6-well plates and cultured for one day and infected with lentiviruses in culture media containing 10 μg/mL protamine sulfate. At one day post-infection, media was replaced with fresh culture media and cells were incubated for two to three additional days to suppress protein expression and to overexpress protein.

Lentivirus production was performed as described ^20^. In brief, HEK293T cells were grown in Dulbecco’s modified eagle’s medium (DMEM) (Gibco/Thermo Fisher Scientific, Gaithersburg, MD, USA) containing 10% FBS (Hyclone, GE healthcare Life Technology, South Logan, UT, USA) and 1% streptomycin/penicillin (Gibco/Thermo Fisher Scientific, Gaithersburg, MD, USA). For lentivirus production, HEK293T cells in 150 mm dishes (Corning Falcon ®, NY, USA) were transfected with 3^rd^ generation lentivirus packaging plasmids, pMDLg/pRRE, pRSV-Rev, and pMD2.G, using TransIT-LT1 transfection reagent (MirusBio, Wisconsin, USA) and incubated for one day, followed by two days of incubation with high serum (30% FBS)-containing DMEM. Lentivirus supernatant was filtered with a 0.45-μm pore filter (EMD Millipore Corp, USA) and concentrated at 20,000 rpm for 2 hrs. Lentiviruses were resuspended in 1 mL in high serum (30% FBS)-containing DMEM and stored at −70°C. To suppress protein expression, HDMVECs grown in 6-well plates were infected with 60 μL of lentiviruses and incubated for 3-4 days, leading to >80% suppression of protein. For overexpression of transgenes, HDMVECs cultured in 6-well plates were infected with 100-200 μL of lentivirus and incubated for 3-4 days. Suppression and overexpression of protein were confirmed with immunoblot or immunofluorescence staining.

### Antibodies and Reagents

Mouse monoclonal anti-human VE-cadherin antibody (clone 55-7H1), polyclonal rabbit anti-human septin 2 (Atlas Antibodies, Cat. # HPA018481), mouse monoclonal anti-human GAPDH (clone 6C5), mouse monoclonal anti-GFP antibody (9F9.F9) were obtained and used as described ^20^. Anti-human nectin-2 antibody (Cat.# AF2229) and anti-human afadin antibody (clone 851204) were obtained from R & D Systems (Minneapolis, MN, USA). Anti-human PECAM-1 antibody (JC/70A) was obtained from Abcam (Cat.# ab9498) (Cambridge, MA, USA) and anti-human ZO-1 antibody was obtained from Invitrogen / ThermoFisher (Carlsbad, CA, USA). Secondary donkey anti-goat IgG conjugated with alexa-568 (Cat.# ab175704), donkey anti-rabbit IgG conjugated with alexa 488 (Cat.# ab150073), and donkey anti-mouse conjugated with alexa 488 (Cat.# ab150105) were obtained from Abcam (Cambridge, MA, USA). Phalloidin conjugated with fluorescent Alexa-647 and secondary antibodies conjugated with horseradish peroxidase were obtained from Molecular Probes ThermoFisher (Eugene, OR, USA) and Sigma-Aldrich (St Louis, MO, USA), respectively.

### Reagents and buffers

Tumor necrosis factor alpha (TNF-α) was obtained from Gibco ThermoFisher (Gaithersburg, MD, USA), and fibronectin was obtained from Sigma-Aldrich (St Louis, MO, USA). ProLong Gold anti-fade solution was obtained from Molecular Probes ThermoFisher (Eugene, OR, USA). T4 DNA Ligase were obtained from Invitrogen ThermoFisher (Carlsbad, CA, USA). PfuTurbo DNA polymerase and QuikChange Site-Directed Mutagenesis Kit were obtained from Agilent Technologies (Santa Clara, CA, USA).

Plasmids expressing two shRNA oligonucleotides targeting septin 2 and one shRNA oligonucleotide targeting LacZ, to serve as a control, were obtained and used as described ^20^.

### Site-directed mutagenesis and Cloning

To create PIP_2_ binding mutant (PIP_2_BM) septin 2, we changed arginine (R) and lysine (K) residues at positions 29 and 30 and lysine residues at positions 33 and 34 to alanine (A) using the QuikChange Site-directed Mutagenesis Kit (Agilent Technologies) according to the manufacturer’s protocol. pBOB-wt septin 2-GFP ^20^ was used as a template for PCR-amplified mutagenesis with primers containing multiple mutation sites. Lentivirus for expression of PIP_2_BM septin 2-GFP was produced as described ^20^.

### Immunofluorescence staining

HDMVEC monolayers were fixed by adding an equal volume of 2X fixation solution composed of freshly dissolved 5% paraformaldehyde in PIPES buffer, prewarmed to 37°C ^57^, directly into culture dishes and incubating at 37°C for 10 min. After removing the fixation solution, cell samples were permeabilized by incubation in 0.1% Triton X-100 in PBS for 5 min at RT. Samples were washed with PBS three times, incubated in blocking buffer (3% BSA in PBS) for at least 30 min, followed by primary antibodies overnight at 4°C, and secondary antibodies conjugated with Alexa fluorescent dyes for 1 hr at RT. Antibodies were diluted in blocking buffer. Samples were washed with PBS three times after primary and secondary antibody incubations. For phalloidin staining, fluorescent phalloidin was added to the secondary antibody solution. Samples were mounted with ProLong Gold anti-fade reagent.

Fluorescence images were collection with a Nikon A1R resonant scanning confocal system using a 40x objective. Z-stacks were collected with a 0.25 μm step size. Images are presented as 2-D projections of Z-stacks prepared with the Fiji implementation of ImageJ open-source software (https://fiji.sc/) ^58^.

### Immunoblots

HDMVECs were washed with RT PBS and lysed by adding 1X SDS-PAGE loading buffer into culture dishes. These whole-cell lysates (WCL) were harvested by scraping the dish and boiling the sample in a heating block for 10 min. WCL was loaded onto 4-20% gradient SDS polyacrylamide gels, separated by electrophoresis, and transferred to PVDF membrane (MilliporeSigma Corp., St Louis, MO, USA) at 90 V for 1 hr at 4°C. The membrane was incubated in blocking buffer (4% BSA in Tris-buffered saline, 0.1% Tween 20 (TBST) buffer) for 30 min at RT and in a primary antibody mixture overnight at 4°C. After washing with PBS with 5 times, the membrane was incubated in horseradish-peroxidase-conjugated secondary antibodies at RT for 1 hr. The membrane was incubated with enhanced chemiluminescence solution for 5 min at RT and developed for autoradiography.

### RNA Sequencing

HDMVECs were grown in 100-mm dishes, treated with 20 ng/mL TNF-α overnight, and harvested for total RNA preparation using RNeasy mini kit from Qiagen (Germantown, MD, USA). Samples were prepared according to manufacturer protocol, indexed, pooled, and sequenced on an Illumina HiSeq system by the Genome Technology Access Center (GTAC) of Washington University School of Medicine. Base calls and demultiplexing were performed with Illumina’s bcl2fastq software and a custom python demultiplexing program with a maximum of one mismatch in the indexing read. RNA-Seq reads were then aligned to the Ensembl release 76 primary assembly with STAR version 2.5.1a ^59^. Gene counts were derived from the number of uniquely aligned unambiguous reads by Subread:featureCount version 1.4.6-p5 ^60^. Isoform expression of known Ensembl transcripts were estimated with Salmon version 0.8.2 ^61^. Sequencing performance was assessed for the total number of aligned reads, total number of uniquely aligned reads, and features detected. The ribosomal fraction, known junction saturation, and read distribution over known gene models were quantified with RSeQC version 2.6.2 ^62^.

All gene counts were then imported into the R/Bioconductor package EdgeR ^63^ and TMM normalization size factors were calculated to adjust for samples for differences in library size. Ribosomal genes and genes not expressed in the smallest group size minus one sample greater than one count-per-million were excluded from further analysis. The TMM size factors and the matrix of counts were then imported into the R/Bioconductor package Limma ^64^. Weighted likelihoods based on the observed mean-variance relationship of every gene and sample were then calculated for all samples with the voomWithQualityWeights ^65^. The performance of all genes was assessed with plots of the residual standard deviation of every gene to their average log-count with a robustly fitted trend line of the residuals. Differential expression analysis was then performed to analyze for differences between conditions and the results were filtered for only those genes with Benjamini-Hochberg false-discovery rate adjusted p-values less than or equal to 0.05.

For each contrast extracted with Limma, global perturbations in known Gene Ontology (GO) terms, MSigDb, and KEGG pathways were detected using the R/Bioconductor package GAGE ^66^ to test for changes in expression of the reported log 2 fold-changes reported by Limma in each term versus the background log 2 fold-changes of all genes found outside the respective term. The R/Bioconductor package heatmap3 ^67^ was used to display heatmaps across groups of samples for each GO or MSigDb term with a Benjamini-Hochberg false-discovery rate adjusted p-value less than or equal to 0.05. Perturbed KEGG pathways where the observed log 2 fold-changes of genes within the term were significantly perturbed in a single-direction versus background or in any direction compared to other genes within a given term with p-values less than or equal to 0.05 were rendered as annotated KEGG graphs with the R/Bioconductor package Pathview ^68^.

## Results

We found previously that septins are required for the proper organization of VE-cadherin at cell junctions of microvascular endothelial monolayers composed of human primary endothelial cells by observing that loss of septin 2 disrupted VE-cadherin structure, membrane dynamics, and junctional integrity ^20^. The junctional integrity of the endothelial monolayer is tightly regulated by a number of diverse intercellular adhesion proteins at cell-cell contact sites in response to cellular and molecular stimuli ^69^. In this study, we extended our analysis by asking whether septin 2 is required to regulate the organization of other adhesion and junctional proteins, including nectin-2, afadin, PECAM-1 and ZO-1.

We localized these junctional proteins in endothelial monolayers of primary human dermal microvascular cells (HDMVECs) in which septin 2 was depleted by shRNA expression. Figure 1 shows immunofluorescence staining for endogenous septin 2 (green), intercellular adhesion proteins (red), and actin filaments, in response to septin 2 suppression. In control cells, nectin-2 (Fig. 1A, upper panels, with inset) and afadin (Fig. 1B, upper panels, with inset) are arranged as continuous thin structures at cell junctions (yellow arrows in insets). In septin 2 suppressed cells, the organization of the proteins changed substantially, to one that was discontinuous, broader and spiky (Fig. 1. A and B. Lower panels). The organization of PECAM-1 (Fig. 1C) also showed dramatic changes, similar to those for nectin-2 and afadin, upon depletion of septin 2. In addition, we examined the effect of septin 2 suppression on a tight junction component, ZO-1 (Fig.1D). ZO-1 also displayed a disorganized arrangement in septin 2 suppressed cells (Fig.1D, lower panels, with Inset). Taken together, septin 2 suppression caused disorganization of all of these various intercellular junctional components of microvascular endothelial cells.

**Figure 1.**
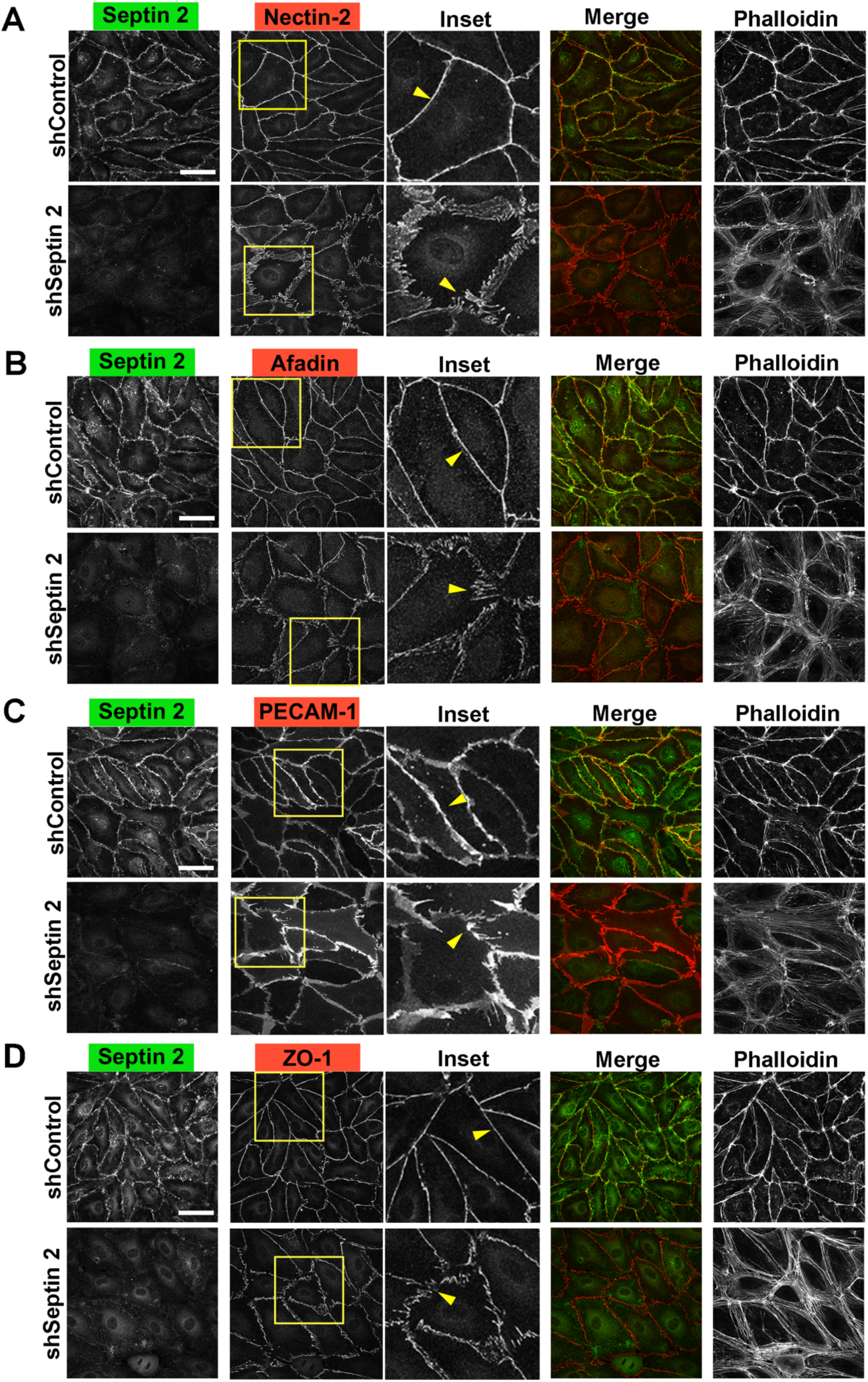
Suppression of septin 2 protein results in disorganization of cell-cell junction proteins. Immunofluorescence staining for endogenous septin 2 (green), cell-cell adhesion proteins (red) and actin filaments in HDMVECs (A-D). (A) Endogenous septin 2, nectin-2, and actin filaments are shown in control (Upper) and septin 2 suppressed HDMVECs (Lower). (B) Endogenous septin 2, afadin, and actin filaments are shown in control (Upper) and septin 2 suppressed HDMVECs (Lower). (C) Endogenous septin 2, PECAM-1, and actin filaments are shown in control (Upper) and septin 2 suppressed HDMVECs (Lower). (D) Endogenous septin 2, tight junction protein (ZO-1), and actin filaments are shown in control (Upper) and septin 2 suppressed HDMVECs (Lower). Inset shows the enlarged area of cell-cell adhesion molecules. Scale bars: 50 μm.

Next, we asked whether the inflammatory mediator TNF-α would elicit disorganization of the same junctional proteins. In our previous work, we found that septin 2 filaments, normally enriched at cell junctions, were depleted from these regions and accumulated in the cytoplasm, in response to TNF-α treatment. VE-cadherin organization was also disrupted, in a manner similar to that of septin 2 depletion. Here, we examined the effect of TNF-α treatment on the arrangement of other intercellular adhesion proteins (Fig. 2). Control untreated HDMVEC monolayers showed the continuous thin arrangement of adhesion proteins described above (Fig. 2, A-D, upper panels with insets). TNF-α treated cells showed dramatically altered arrangements of nectin-2, afadin, PECAM-1, and ZO-1 (Fig. 2, A-D, lower panels, with insets). In addition, TNF-α treatment led to loss of septin 2 from near cell junctions with accumulation in the cytoplasm.

**Figure 2.**
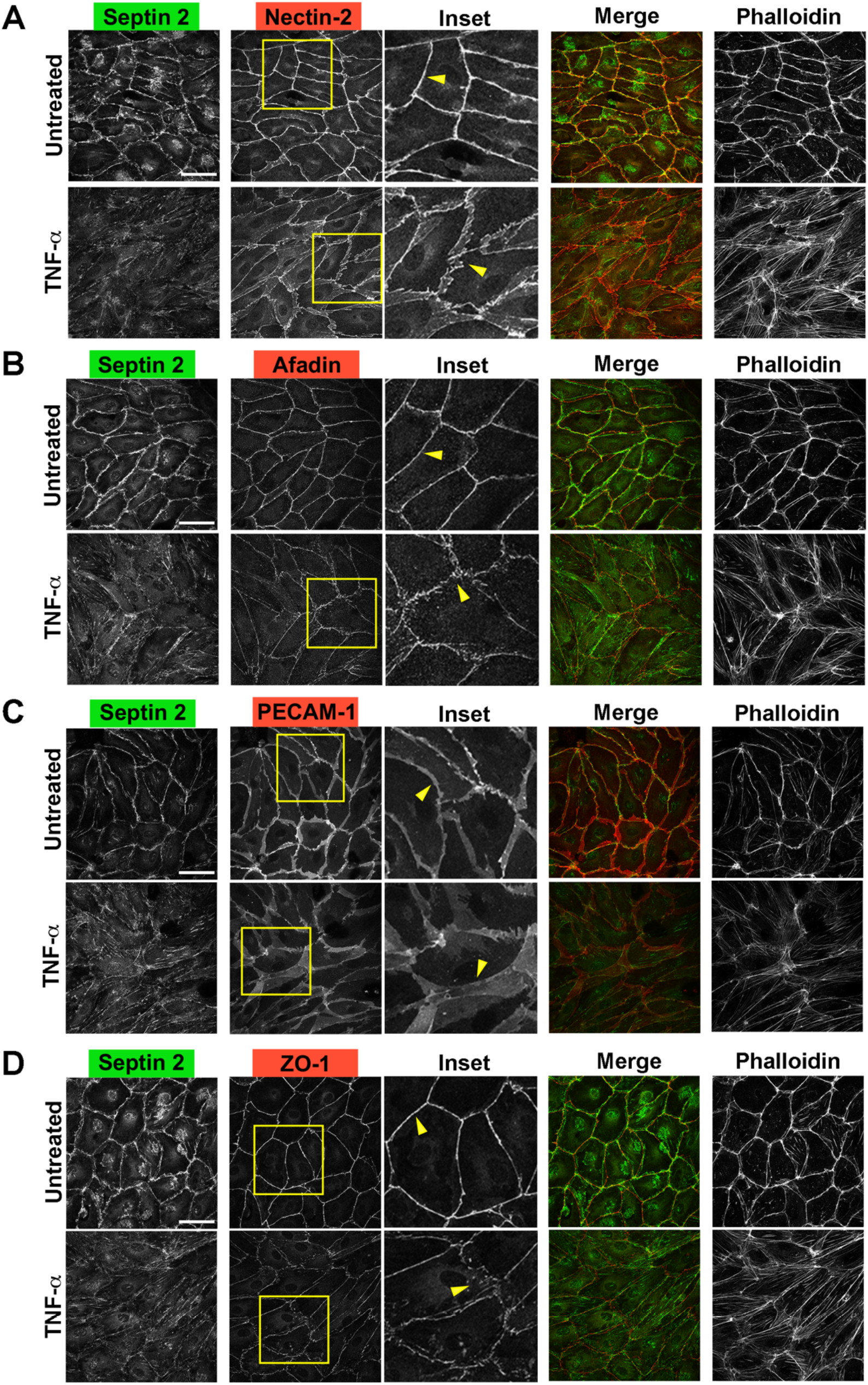
TNF-α treatment sequesters septin 2 to the cytoplasm in association with disorganization of cell-cell junction proteins. Immunofluorescence staining for endogenous septin 2 (green), cell-cell adhesion proteins (red) and actin filaments (A-D). (A) Endogenous septin 2, nectin-2, and actin filaments are shown in untreated (Upper) and TNF-α-treated HDMVECs (Lower). Inset indicates enlarged nectin-2 structure at the junctions. (B) Endogenous septin 2, afadin, and actin filaments are shown in untreated (Upper) and TNF-α-treated HDMVECs (Lower). Inset indicates enlarged afadin structure at the junctions. (C) Endogenous septin 2, PECAM-1, and actin filaments are shown in untreated (Upper) and TNF-α-treated HDMVECs (Lower). Inset indicates enlarged PECAM-1 structure at the junctions. (D) Endogenous septin 2, ZO-1, and actin filaments are shown in untreated (Upper) and TNF-α-treated HDMVECs (Lower). Inset indicates enlarged ZO-1 structure at the junctions. Scale bars: 50 μm.

We hypothesized that septin 2 localizes to the plasma membrane at areas of positive curvature via association of its polybasic domain with acidic phospholipids. Previous studies found that arginine and lysine residues in the polybasic domain of human septins are important for binding to PIP_2_ and PIP_3_ *in vitro* and in cells and that these basic residues are conserved in the polybasic domain of septin 2 ^12,13,70–72^. Therefore, we asked whether mutation of basic residues in the polybasic domain of septin 2 would disrupt septin 2 accumulation and organization of endothelial cell junction proteins. We mutated arginine (R) and lysine (K) residues to alanine (A) in the poly basic domain at the N-terminal region of septin 2, as diagrammed in Figure 3 A. We expressed this PIP_2_ binding mutant (PIP_2_ BM) form of septin 2 in HDMVECs using a pBOB-GFP lentiviral expression system. We compared expression of wild-type (wt) septin 2 fused with GFP with expression of PIP_2_BM septin 2 fused with GFP. Using immunofluorescence staining, we imaged the distribution of expressed septin 2-GFP with antibodies to GFP and the distribution of total septin 2 (endogenous and overexpressed) with antibodies to septin 2 (Supplemental Fig. 1). Expressed wt septin 2 (green) localized to cell junctions (Supplemental Fig. 1A, cells with blue pentagons), and its expression did not change the localization of endogenous septin 2 at the junctions (cells with blue pentagons versus cells will magenta dots). In contrast, cells expressing PIP_2_BM mutant septin 2-GFP showed a dominant effect, with altered localization of septin 2, consisting of loss of endogenous septin 2 (red) from junctional localizations, with accumulation in the cytoplasm (Supplemental Fig. 1B, Inset, cells with blue pentagons), by comparison with nearby cells that were not infected and did not express PIP_2_ BM septin 2 (Supplemental Fig. 1B, Inset, cells with magenta dots). Thus, expression of PIP_2_BM septin 2 had a dominant-negative effect on the distribution and localization of endogenous wt septin 2.

**Figure 3.**
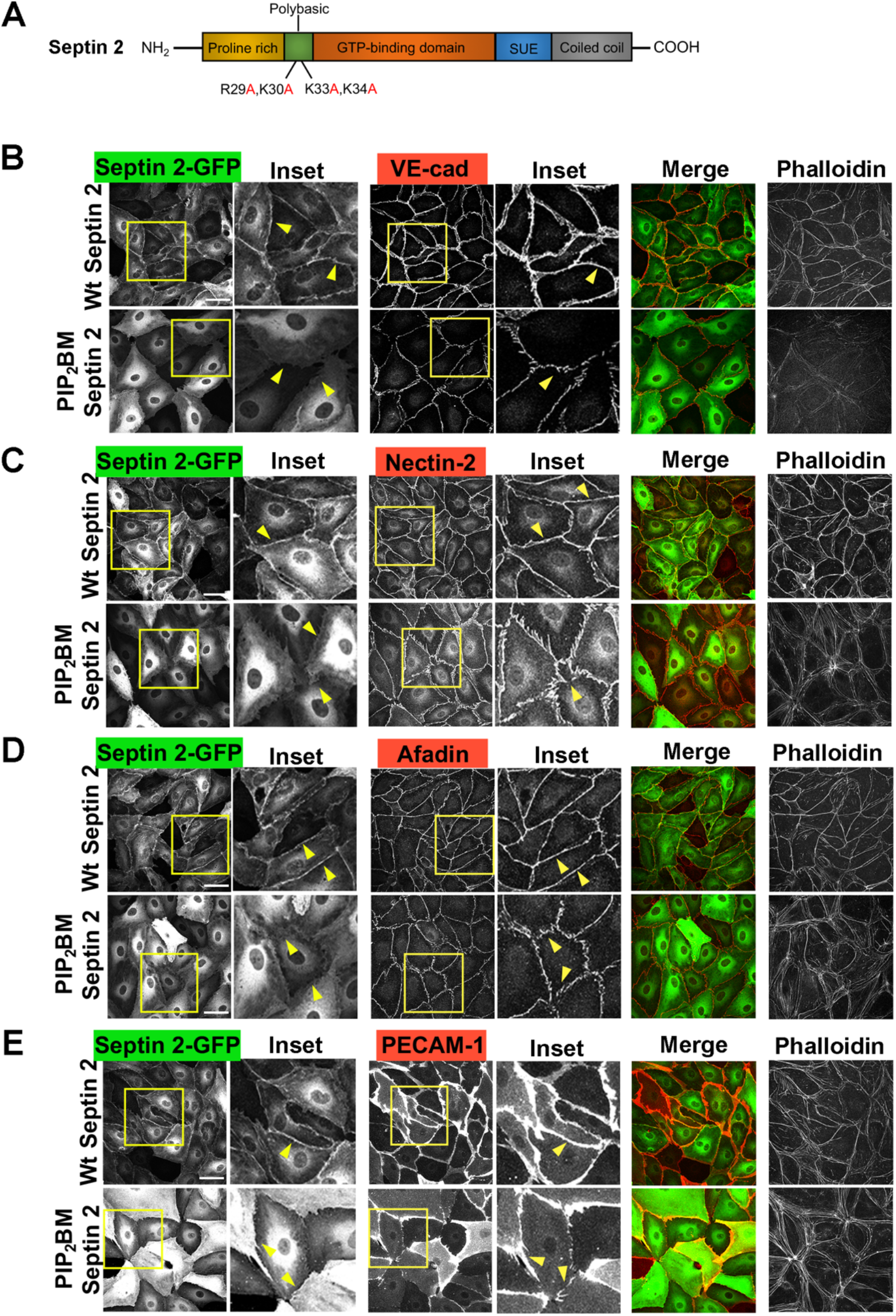
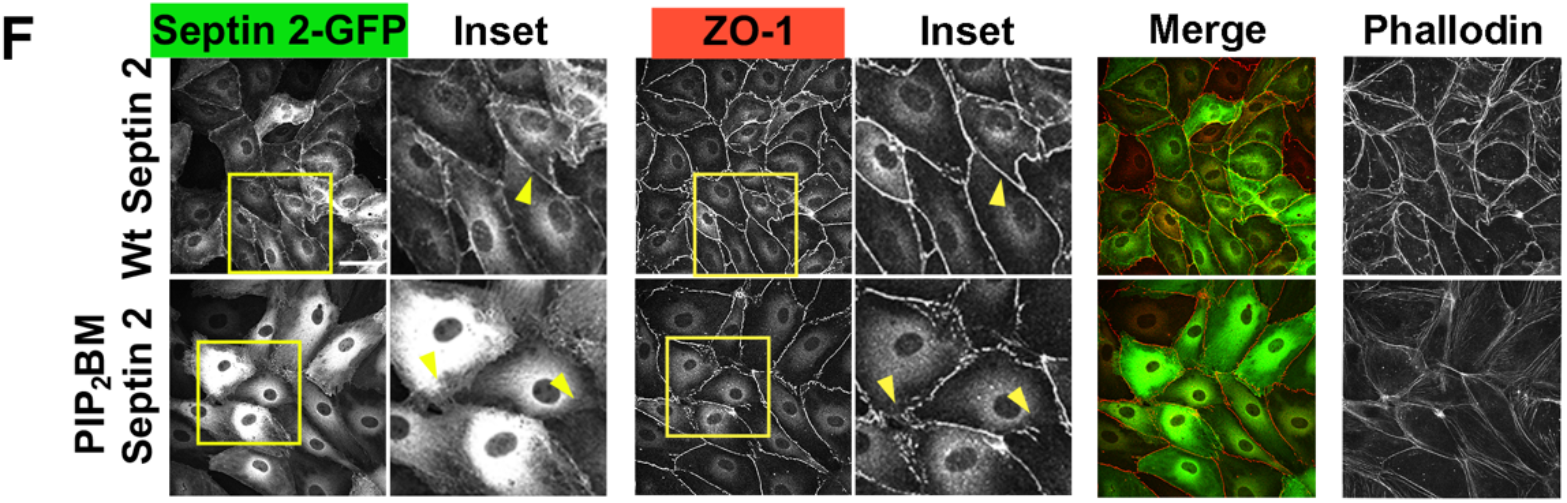
Expressed polybasic mutant septin 2 fails to localize to the junction. (A) Diagram of septin 2 domain architecture shows basic residues, R and K in the polybasic domain at the N-terminus. (B-E) Immunofluorescence staining shows overexpression of wild type (wt) and PIP_2_ binding mutant (BM) septin 2 (green), cell-cell junction protein (red), and actin filaments. (B) Overexpression of wt septin 2-GFP and PIP_2_BM septin 2-GFP (Upper, Inset) and organization of VE-cadherin at the junctions (Lower, Inset). (C) Overexpression of wt septin 2-GFP and PIP_2_BM septin 2-GFP (Upper, Inset) and organization of nectin-2 at the junctions (Lower, Inset). (D) Overexpression of wt septin 2-GFP and PIP_2_BM septin 2-GFP (Upper, Inset) and organization of afadin at the junctions (Lower, Inset). (E) Overexpression of wt Septin 2-GFP and PIP_2_BM Septin 2-GFP (Upper, Inset) and organization of PECAM-1 at the junctions (Lower, Inset). Overexpression of wt septin 2-GFP and PIP_2_BM septin 2-GFP (Upper, Inset) and organization of ZO-1 at the junctions (Lower, Inset). Scale bars: 50 μm.

We asked whether expression of PIP_2_BM septin 2 affected the arrangement and organization of endothelial junctional proteins by immunofluorescence staining of cells expressing PIP_2_BM septin 2 versus wt septin 2 (Fig. 3, B-F). First, we found that wt septin 2 expression did not affect the continuous thin-line arrangement of VE-cadherin at cell junctions (Fig. 3B, upper panels, with inset). On the other hand, expression of PIP_2_BM septin 2 dramatically disrupted the organization of VE-cadherin structures at cell junctions (Fig. 3B, lower panels, with inset), in a manner similar that produced by septin 2 suppression and by TNF-α treatment ^20^. Similar effects were observed for other junctional proteins, including nectin-2 (Fig. 3C), afadin (Fig. 3D), and PECAM-1 (Fig. 3E) as well as the tight junction protein ZO-1 (Fig. 3F). Again, these effects were similar to the effects of septin 2 depletion and TNF-α treatment (Figure 1 and 2). We conclude that septin 2 tethering to the plasma membrane via interaction of arginine and lysine residues in the polybasic domain of septin 2 with anionic phosphoinositides of the plasma membrane is required for the proper organization and integrity of junctional proteins.

We considered the possibility that the effects of septin 2 suppression and TNF-α treatment might be mediated by effects on the level of expression of the various junction-associated proteins. We reasoned that cooperative assembly of filament and macromolecular complexes might be affected in a non-linear manner by decreases in levels of protein components. We examined expression levels in two ways – by protein immunoblots with specific antibodies and by RNA-Seq analysis of whole-cell RNA samples. Immunoblot analysis revealed that the nectin-2 protein level was increased in response to septin 2 suppression (Fig. 4A) and TNF-α treatment (Fig. 4B). The protein levels of afadin, PECAM-1 and ZO-1 did not change significantly with either septin 2 depletion (Fig. 4A) or with TNF-α treatment (Fig. 4B). As noted above, the effects of septin 2 suppression and TNF-α treatment were similar to each other.

**Figure 4.**
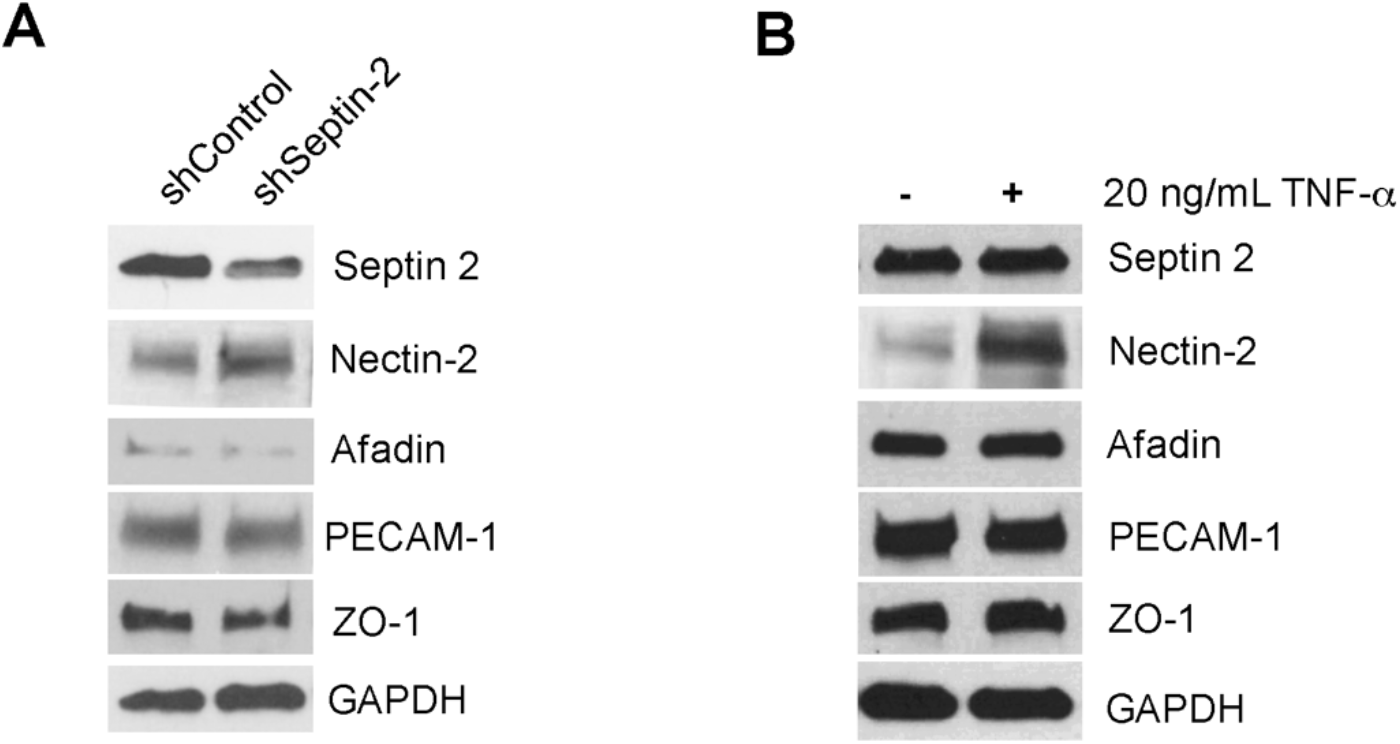
Expression levels of intercellular adhesion proteins in response to septin 2 depletion and TNF-α treatment. Immunoblot analysis for junctional adhesion proteins. Expression levels of each protein are shown in response to (A) septin 2 depletion and (B) TNF-α treatment. GAPDH is an internal loading control.

We also performed RNA-Seq analysis of whole-cell RNA samples from HDMVECs to examine expression of the genes encoding cell junction proteins at the RNA level (Fig. 5 and Supplemental table 1-4). Differential gene expression profiles in response to septin 2 suppression and TNF-α treatment are shown in Volcano plots (Fig. 5), and genes with significant expression changes are listed in Supplemental Tables 1 and 2. Examining the results for all genes, we found significant levels of up- and down-regulation (over log_2_ fold change, log_2_FC), as indicated by blue and red color, respectively (Fig. 5). For septin 2 suppression, the genes ADAM9, ACER2, CYGB, and ENKUR were among the ones up-regulated most strongly, while FLN4, ENTPD5, PCDDH10, NDRG4, COL1A2, and CNR1 were among those down-regulated most strongly (Fig. 5A). Treatment with TNF-α led to significant changes in a different set of genes, with strongest up-regulation of U2AF1, CXCL11, BEST1 and SPARCL1, and strongest down-regulation of SPATA13, FAAH, CCL1, WHRN, FOXF1, and PHOD3 (Fig. 5B). Lists of genes that change expression more than two-fold in response to septin 2 suppression and TNF-α treatment are available in supplemental table 1 and 2.

**Figure 5.**
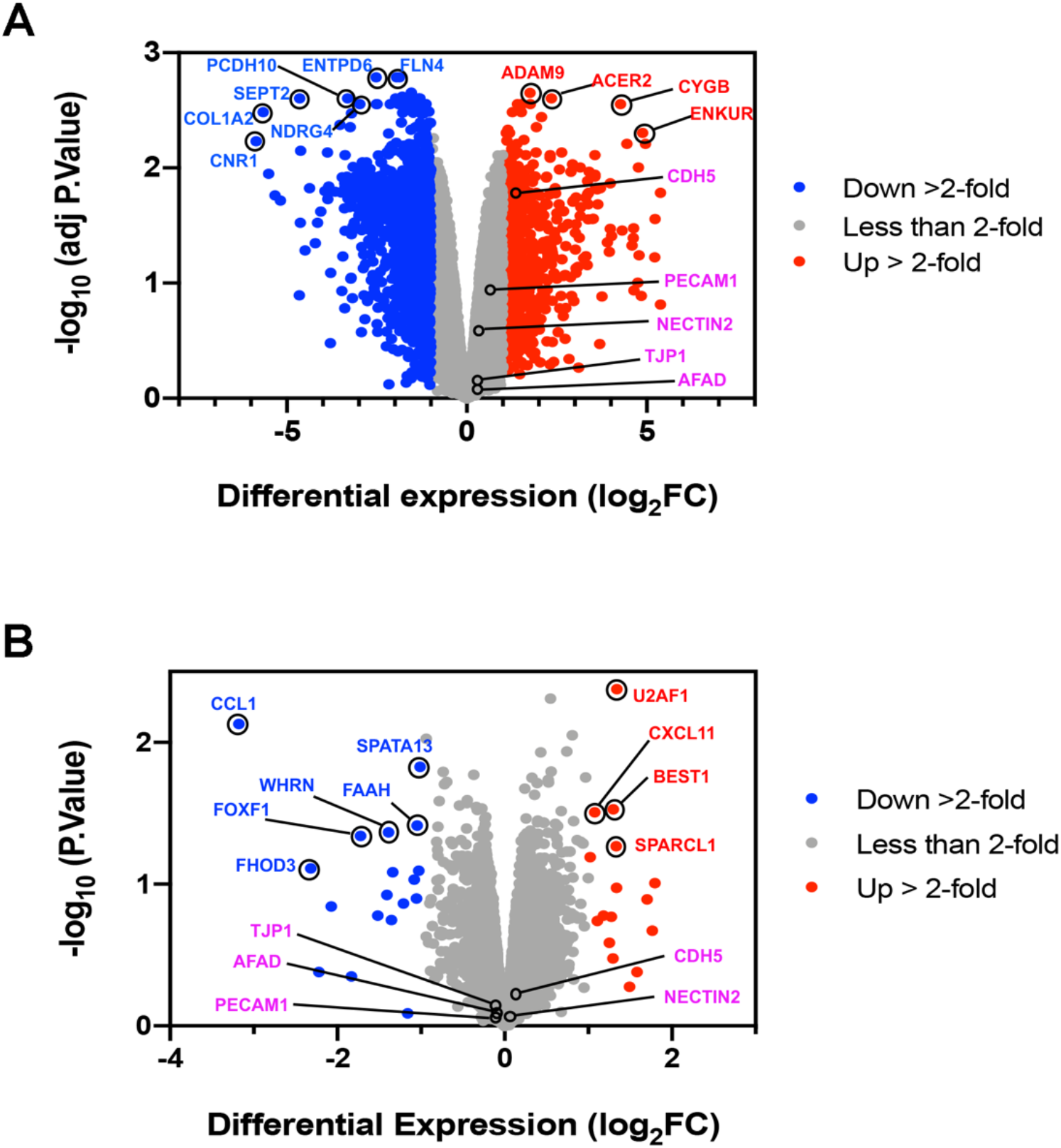
Differential gene expression profiles in septin 2 suppressed and TNF-α treated HDMVECs. Volcano plot shows differential gene expression profile from RNA sequence with (A) septin 2-suppressed HDMVECs and (B) TNF-α treated HDMVECs. Up-regulated genes (red) and down-regulated genes (blue) are shown. Blue and red coloring indicates gene expression changes over 1 of log_2_FC (fold change).

We looked specifically at RNA levels for genes encoding the junctional proteins examined above, namely VE-cadherin (CDH5), nectin-2 (NECTIN2), afadin (AFND), PECAM-1 (PECAM1), and ZO-1 (TJP1). VE-cadherin (CDH5) RNA levels were significantly increased by septin 2 suppression and slightly increased by TNF-α treatment (Fig, 5A and B, pink), consistent with the protein expression analysis by immunoblot in our previous study ^20^. We observed no significant changes in expression for nectin-2, afadin, PECAM-1, or ZO-1 (Fig. 5A and B, pink, Supplemental Tables 3 and 4).

## Discussion

### Endothelial Cell Junction Components and Septins

In this study, we discovered that junctional localization of septin 2 is required to preserve the proper organization of a number of cell junctional proteins of microvascular endothelial cell monolayers. We focused on the adherens junction proteins, nectin-2 and afadin, the tight junction protein ZO-1, and the general adhesion protein PECAM-1, extending our previous study of VE-cadherin ^20^. We found striking disorganization of all these junctional proteins in response to depletion of septin 2 protein.

We considered that this disorganization of the junctional proteins might result from decreased expression levels of the proteins, especially if their assembly involved positive cooperativity. However, we found no changes in the levels of afadin, PECAM-1, and ZO-1, and we found increases in protein level for nectin-2. Thus, the disruption of their junctional organization was not due simply to a loss of protein. These results are similar to those observed for VE-cadherin in our previous study ^20^. Our results for gene expression, based on RNA levels by RNA-Seq, are consistent with the results from assays of protein level.

The most likely role for septins in the organization of these various junctional components, in our view, is to support the positively curved regions of membrane at the base of actin-based lamellipodial protrusions, which was our conclusion in our previous study ^20^. We would suggest that endothelial cells are constantly forming protrusions that push toward each other to form adhesions, as indicated by previous studies ^54,55^, most notably that of Efimova and Svitkina ^73^. The accumulated evidence supports a model in which sites of cell-cell contact, where membrane protrusions from apposing cells meet, are locations where adhesive junctional cell surface molecules interact with their counterparts on adjacent cells and assemble into the supramolecular structures defined as cell junctions.

The junctional components that we examined included ones found in endothelial cell adherens junctions, in tight junctions and diffusely located on the surface. Adherens junctions of endothelial cells contain VE-cadherin, nectin-2 and afadin ^74^. Nectin-2 and afadin are recruited to VE-cadherin-containing adherens junctions in both epithelial cells ^75^ and endothelial cells ^76^. PECAM-1 is localized at intercellular adhesion sites between endothelial cells and is also found at contacts between endothelial cells and platelets ^77^. Tight junctions of endothelial cells include the protein Zona Occludens-1 (ZO-1).

### Septin - Membrane Interactions

Septin proteins and filaments are known to interact directly with membrane lipids ^13,70–72,78^, accumulating at locations with positive curvature ^18,79,80^. This direct interaction can be mediated by a polybasic region of septins interacting with acidic head groups of phospholipids on the membrane surface ^81^. To investigate whether an electrostatic interaction mechanism might also account for the association of septins with plasma membranes in endothelial cells, we tested the effects of mutating the polybasic region of septin 2, substituting basic residues with neutral ones. Indeed, as predicted, the mutant septin 2 failed to localize to membranes. Moreover, expression of the mutant septin 2 had a dominant effect, revealed by the loss of endogenous septin 2 from the membrane. This dominant effect is reasonable because septins form hetero-oligomers that assemble into filaments. The partial loss of the polybasic septin / acidic membrane lipid interaction is sufficient to cause the loss of septins from the membrane, consistent with additive interactions being necessary for association.

### TNF-α and Septins

As part of the process of inflammation, the cytokine TNF-α causes increased permeability of the endothelium with partial loss of cell junction integrity ^20^. We asked whether septin 2, based on its role in cell junction integrity, might be a component of the mechanism of action of TNF-α^55,82^. Indeed, we found that TNF-α treatment of endothelial monolayers causes loss of septin 2 filaments normally enriched at cell junctions, with shifting of their distribution to the cytoplasm. This change in septin 2 distribution would appear to be sufficient for the disorganization of junctional proteins that results from TNF-α treatment.

One might speculate that the membrane tethering of septin 2 by anionic phospholipids would be regulated by TNF-α induced signaling pathways. TNF-α has been shown to activate PLC γ, which catalyzes the hydrolysis of PIP_2_ at the membrane into IP_3_ and DAG ^83^. Thus, TNF-α may induce decreases in PIP_2_ levels at the membrane. This speculative hypothesis is likely to be an oversimplification. Treatment with TNF-α has many effects in addition to the effects on septin 2 localization. For example, our gene expression analysis showed many notable differences between septin 2 depletion and TNF-α treatment,

### Potential Cooperative Role for Actin Filaments

Septin 2 suppression, PIP_2_BM septin 2 expression, and TNF-α treatment all resulted in loss of F-actin from accumulations at the membrane, in association with cell junctions, into the cytoplasm. Septin filament networks and actin filament networks show functional interdependence ^26–31^. Therefore, the role of septin filaments at cell junctions may involve cooperative interactions among cytosolic actin filaments and cell junction.

### Summary

Our results are summarized in a schematic diagram in Figure 6, which illustrates intercellular adhesion proteins and septin 2 at the junctions of endothelial monolayers. Under basal conditions, septin 2 is localized at the junctions mediated by interaction between PIP_2_ at the membrane and basic residues in septin 2. The cell junctions are associated with well-organized structures of intercellular adhesion proteins, such as VE-cadherin, nectin-2, afadin, PECAM-1, and ZO-1. Septin 2 is enriched at the positively curved areas of membrane (Fig. 6. A). Loss of septin 2 at cell junctions, caused by suppression of septin 2 expression, treatment with TNF-α, and loss of interaction with membrane PIP_2_ causes significant alteration of the arrangement of junctional proteins and defects in the integrity of the barrier formed by endothelial monolayers (Fig. 6. B).

**Figure 6.**
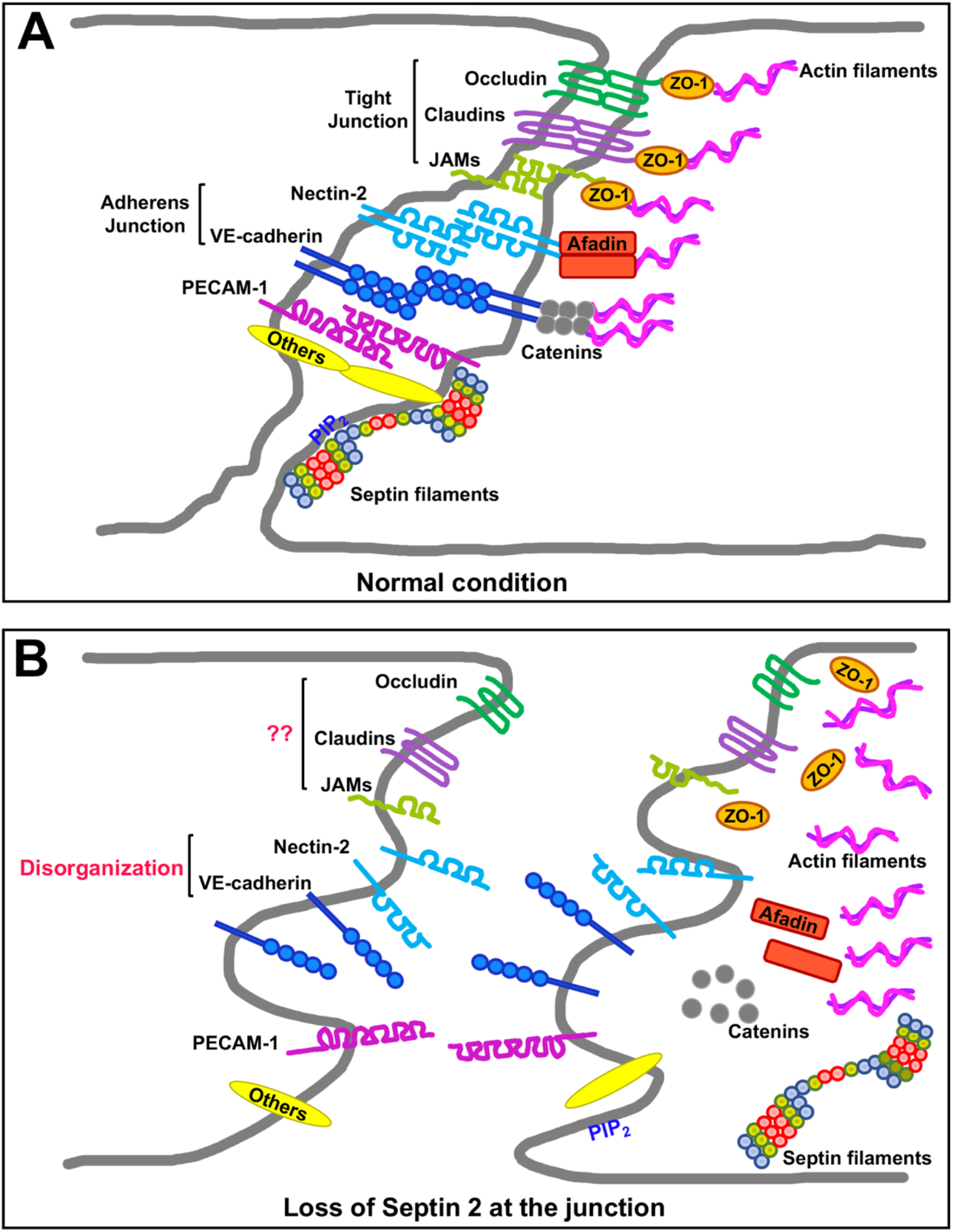
Diagram of septin 2 and intercellular junctional proteins at endothelial cell junctions. (A) Under basal resting conditions, septin 2 filaments are localized at the junctions of microvascular endothelial cells, and various junctional proteins are properly organized at the junctions. (B) Loss of septin 2 at the cells junctions leads to disorganization of junctional components of microvascular endothelial monolayers resulting in loss of junctional integrity.

## Abbreviations

AJ: Adherens junction
BSA: Bovine serum albumin
EGM-2MV: Endothelial cell growth medium-2 microvascular
F-actin: actin filaments
FBS: Fetal bovine serum
GFP: Green fluorescent protein
GJ: Gap junction
HDMVECs: Human dermal microvascular endothelial cells
IgG: Immunoglobulin G
JAM: junctional adhesion molecule
PBS: Phosphate buffered saline
PECAM-1: Platelet-endothelial cell adhesion molecule-1
PIP2: Phosphatidylinositol 4,5-bisphosphate
TBS: Tris buffered saline
TJ: Tight junction
TNF-α: Tumor necrosis factor alpha
VE-cadherin: Vascular endothelial-cadherin
WCL: Whole cell lysate
ZO-1: Zonula occludens-1

## Acknowledgements

We are grateful to members of our laboratory and department for advice and assistance. The Genome Technology Access Center (GTAC) of Washington University School of Medicine provided advice and assistance with RNA sequencing. Light microscopy was performed in part through the use of the Washington University Center for Cellular Imaging (WUCCI) supported by the Washington University School of Medicine, The Children’s Discovery Institute of Washington University, and St. Louis Children’s Hospital (CDI, CORE-2015-505) and NIH / NINDS NS086741.

## Data availability

Raw data for RNA-Seq experiments is available from the authors by direct request and via their laboratory web site (cooperlab.wustl.edu).

---

**Supplemental Figure 1.**
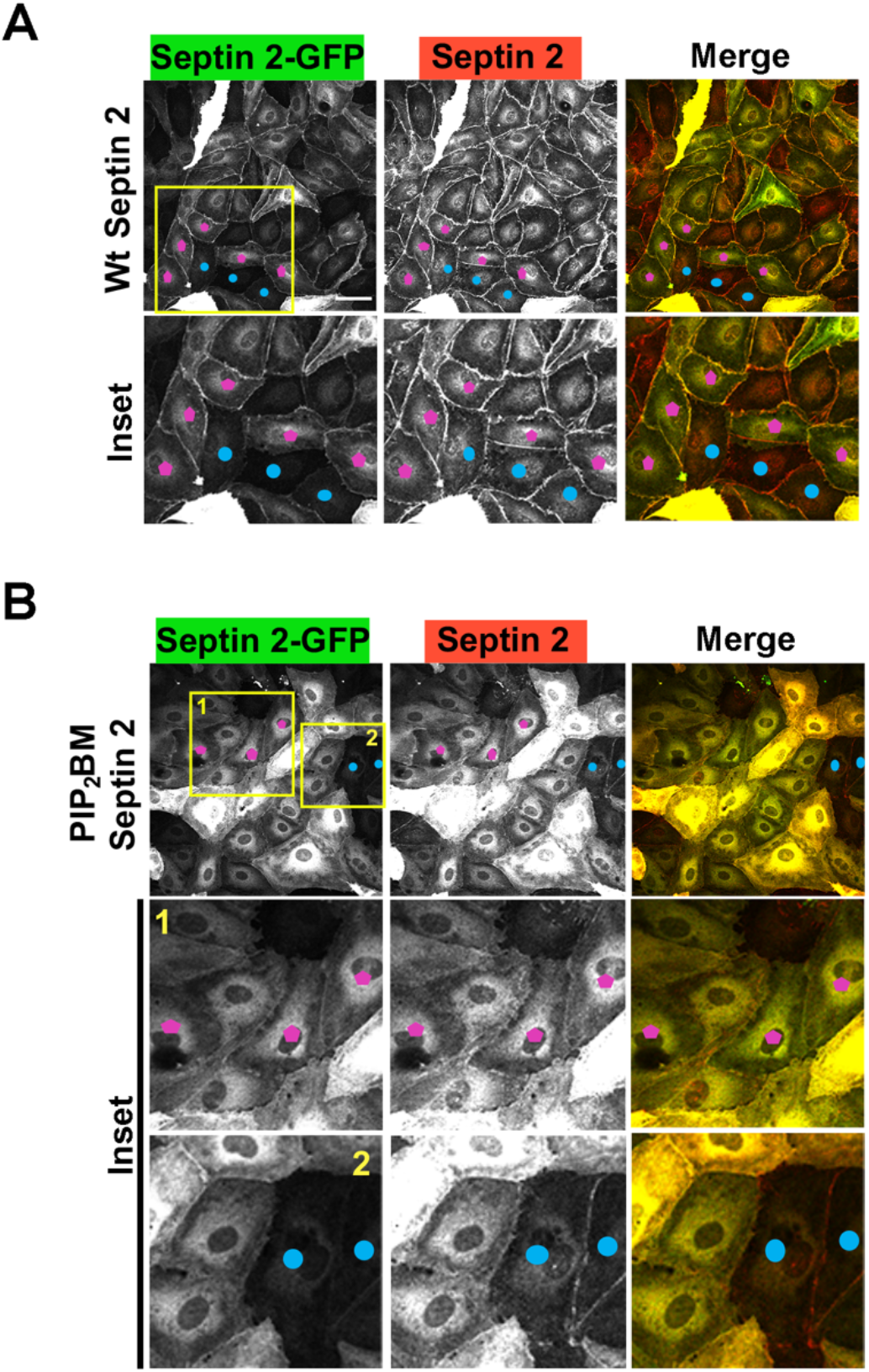
Overexpression of PIP_2_ BM septin 2 shows a dominant negative effect. (A) Immunofluorescence staining for overexpressed wt septin 2-GFP with anti-GFP antibody (green) and total septin 2 (red) with anti-septin 2 antibody. (B) Immunofluorescence staining for overexpressed PIP_2_BM septin 2-GFP with anti-GFP antibody (green) and total septin 2 (red) with anti-septin 2 antibody. Blue pentagons indicate the cells overexpressing septin 2 (wt and PIP_2_BM). Magenta dots indicate non-overexpressing cells. Scale bars: 50 μm.

**Supplemental table 1. List of genes that show significant change in expression following septin 2 suppression.**

**Supplemental table 2. List of genes that show significant change in expression following TNF-α treatment.**

**Supplemental table 3. List of cell adhesion molecule genes that show different expression following septin 2 suppression.**

**Supplemental table 4. List of cell adhesion molecule genes that show different expression by TNF-α treatment.**

## References

1. Hartwell, L. H. Genetic control of the cell division cycle in yeast. IV. Genes controlling bud emergence and cytokinesis. Exp Cell Res. 1971;69:265–276.

2. Haarer, B. K. & Pringle, J. R. Immunofluorescence localization of the *Saccharomyces cerevisiae CDC12* gene product to the vicinity of the 10-nm filaments in the mother-bud neck. Mol. Cell. Biol. 1987;7:3678–3687.

3. Ong, K., Wloka, C., Okada, S., Svitkina, T. & Bi, E. Architecture and dynamic remodelling of the septin cytoskeleton during the cell cycle. Nat Commun. 2014;5:5698.

4. Oh, Y. & Bi, E. Septin structure and function in yeast and beyond. Trends Cell Biol. 2011;21:141–148.

5. Mostowy, S. & Cossart, P. Septins: the fourth component of the cytoskeleton. Nat. Rev. Mol. Cell Biol. 2012;13:183–194.

6. Dolat, L., Hu, Q. & Spiliotis, E. T. Septin functions in organ system physiology and pathology. Biol. Chem. 2014;395:123–141.

7. Kinoshita, M. Assembly of mammalian septins. J. Biochem. 2003;134:491–496.

8. Pan, F., Malmberg, R. L. & Momany, M. Analysis of septins across kingdoms reveals orthology and new motifs. BMC Evol Biol. 2007;7:103.

9. Cao, L. et al. Phylogenetic and evolutionary analysis of the septin protein family in metazoan. FEBS Lett. 2007;581:5526–5532.

10. Bridges, A. A. & Gladfelter, A. S. Septin Form and Function at the Cell Cortex. J. Biol. Chem. 2015;290:17173–17180.

11. Valadares, N. F., d’ Muniz Pereira, H., Ulian Araujo, A. P. & Garratt, R. C. Septin structure and filament assembly. Biophys Rev. 2017;9:481–500.

12. Zhang, J. et al. Phosphatidylinositol polyphosphate binding to the mammalian septin H5 is modulated by GTP. Curr. Biol. 1999;9:1458–1467.

13. Omrane, M. et al. Septin 9 has Two Polybasic Domains Critical to Septin Filament Assembly and Golgi Integrity. iScience. 2019;13:138–153.

14. Bridges, A. A. et al. Septin assemblies form by diffusion-driven annealing on membranes. Proc. Natl. Acad. Sci. U S A. 2014;111:2146–2151.

15. Bertin, A. et al. Three-dimensional ultrastructure of the septin filament network in *Saccharomyces cerevisiae*. Mol. Biol. Cell. 2012;23:423–432.

16. Rodal, A. A., Kozubowski, L., Goode, B. L., Drubin, D. G. & Hartwig, J. H. Actin and septin ultrastructures at the budding yeast cell cortex. Mol. Biol. Cell. 2005;16:372–384.

17. Spiliotis, E. T. & Gladfelter, A. S. Spatial guidance of cell asymmetry: septin GTPases show the way. Traffic. 2012;13:195–203.

18. Bridges, A. A., Jentzsch, M. S., Oakes, P. W., Occhipinti, P. & Gladfelter, A. S. Micron-scale plasma membrane curvature is recognized by the septin cytoskeleton. J. Cell Biol. 2016;213:23–32.

19. Kang, H. & Lew, D. J. How do cells know what shape they are? Curr. Genet. 2017;63:75–77.

20. Kim, J. & Cooper, J. A. Septins regulate junctional integrity of endothelial monolayers. Mol. Biol. Cell. 2018;29:1693–1703.

21. Dolat, L. et al. Septins promote stress fiber-mediated maturation of focal adhesions and renal epithelial motility. J. Cell Biol. 2014;207:225–235.

22. Fung, K. Y., Dai, L. & Trimble, W. S. Cell and molecular biology of septins. Int Rev Cell Mol Biol. 2014;310:289–339.

23. Spiliotis, E. T. & Nelson, W. J. Here come the septins: novel polymers that coordinate intracellular functions and organization. J. Cell Sci. 2006;119:4–10.

24. Pagliuso, A. et al. A role for septin 2 in Drp1-mediated mitochondrial fission. EMBO Rep. 2016;17:858–873.

25. Dolat, L. & Spiliotis, E. T. Septins promote macropinosome maturation and traffic to the lysosome by facilitating membrane fusion. J. Cell Biol. 2016;214:517–527.

26. Bezanilla, M., Gladfelter, A. S., Kovar, D. R. & Lee, W. L. Cytoskeletal dynamics: a view from the membrane. J. Cell Biol. 2015;209:329–337.

27. Hu, J. et al. Septin-driven coordination of actin and microtubule remodeling regulates the collateral branching of axons. Curr. Biol. 2012;22:1109–1115.

28. Joo, E., Surka, M. C. & Trimble, W. S. Mammalian SEPT2 is required for scaffolding nonmuscle myosin II and its kinases. Dev Cell. 2007;13:677–690.

29. Mavrakis, M. et al. Septins promote F-actin ring formation by crosslinking actin filaments into curved bundles. Nat. Cell Biol. 2014;16:322–334.

30. Spiliotis, E. T. Spatial effects - site-specific regulation of actin and microtubule organization by septin GTPases. J. Cell Sci. 2018;;131

31. Spiliotis, E. T., Hunt, S. J., Hu, Q., Kinoshita, M. & Nelson, W. J. Epithelial polarity requires septin coupling of vesicle transport to polyglutamylated microtubules. J. Cell Biol. 2008;180:295–303.

32. Baldwin, A. L. & Thurston, G. Mechanics of endothelial cell architecture and vascular permeability. Crit Rev Biomed Eng. 2001;29:247–278.

33. Deanfield, J. E., Halcox, J. P. & Rabelink, T. J. Endothelial function and dysfunction: testing and clinical relevance. Circulation. 2007;115:1285–1295.

34. Michiels, C. Endothelial cell functions. J. Cell. Physiol. 2003;196:430–443.

35. Sumpio, B. E., Riley, J. T. & Dardik, A. Cells in focus: endothelial cell. Int J Biochem Cell Biol. 2002;34:1508–1512.

36. Bazzoni, G. & Dejana, E. Endothelial cell-to-cell junctions: molecular organization and role in vascular homeostasis. Physiol. Rev. 2004;84:869–901.

37. Dejana, E. Endothelial cell-cell junctions: happy together. Nat. Rev. Mol. Cell Biol. 2004;5:261–270.

38. Lampugnani, M. G., Dejana, E. & Giampietro, C. Vascular Endothelial (VE)-Cadherin, Endothelial Adherens Junctions, and Vascular Disease. Cold Spring Harb Perspect Biol. 2017;doi: 10.1101/cshperspect.a029322.

39. Giannotta, M., Trani, M. & Dejana, E. VE-cadherin and endothelial adherens junctions: active guardians of vascular integrity. Dev Cell. 2013;26:441–454.

40. Sakakibara, S., Maruo, T., Miyata, M., Mizutani, K. & Takai, Y. Requirement of the F-actin-binding activity of l-afadin for enhancing the formation of adherens and tight junctions. Genes Cells. 2018;23:185–199.

41. Tabariès, S. et al. Afadin cooperates with Claudin-2 to promote breast cancer metastasis. Genes Dev. 2019;33:180–193.

42. Heinemann, U. & Schuetz, A. Structural Features of Tight-Junction Proteins. Int J Mol Sci. 2019;;20

43. Buckley, A. & Turner, J. R. Cell Biology of Tight Junction Barrier Regulation and Mucosal Disease. Cold Spring Harb Perspect Biol. 2018;;10

44. McNeil, E., Capaldo, C. T. & Macara, I. G. Zonula occludens-1 function in the assembly of tight junctions in Madin-Darby canine kidney epithelial cells. Mol. Biol. Cell. 2006;17:1922–1932.

45. Tornavaca, O. et al. ZO-1 controls endothelial adherens junctions, cell-cell tension, angiogenesis, and barrier formation. J. Cell Biol. 2015;208:821–838.

46. Ebnet, K., Suzuki, A., Ohno, S. & Vestweber, D. Junctional adhesion molecules (JAMs): more molecules with dual functions. J. Cell Sci. 2004;117:19–29.

47. Kumar, N. M. & Gilula, N. B. The gap junction communication channel. Cell. 1996;84:381–388.

48. Saez, J. C., Berthoud, V. M., Branes, M. C., Martinez, A. D. & Beyer, E. C. Plasma membrane channels formed by connexins: their regulation and functions. Physiol. Rev. 2003;83:1359–1400.

49. Privratsky, J. R. & Newman, P. J. PECAM-1: regulator of endothelial junctional integrity. Cell Tissue Res. 2014;355:607–619.

50. Hahn, J. H. et al. CD99 (MIC2) regulates the LFA-1/ICAM-1-mediated adhesion of lymphocytes, and its gene encodes both positive and negative regulators of cellular adhesion. J. Immunol. 1997;159:2250–2258.

51. Schenkel, A. R., Mamdouh, Z., Chen, X., Liebman, R. M. & Muller, W. A. CD99 plays a major role in the migration of monocytes through endothelial junctions. Nat. Immunol. 2002;3:143–150.

52. Hartsock, A. & Nelson, W. J. Adherens and tight junctions: structure, function and connections to the actin cytoskeleton. Biochim. Biophys. Acta. 2008;1778:660–669.

53. Garcia-Ponce, A., Citalán-Madrid, A. F., Velázquez-Avila, M., Vargas-Robles, H. & Schnoor, M. The role of actin-binding proteins in the control of endothelial barrier integrity. Thromb. Haemost. 2015;113:20–36.

54. Mooren, O. L., Kim, J., Li, J. & Cooper, J. A. Role of N-WASP in Endothelial Monolayer Formation and Integrity. J. Biol. Chem. 2015;290:18796–18805.

55. Mooren, O. L., Li, J., Nawas, J. & Cooper, J. A. Endothelial cells use dynamic actin to facilitate lymphocyte transendothelial migration and maintain the monolayer barrier. Mol. Biol. Cell. 2014;25:4115–4129.

56. Schnittler, H. et al. Actin filament dynamics and endothelial cell junctions: the Ying and Yang between stabilization and motion. Cell Tissue Res. 2014;355:529–543.

57. Mooren, O. L., Kotova, T. I., Moore, A. J. & Schafer, D. A. Dynamin2 GTPase and cortactin remodel actin filaments. J. Biol. Chem. 2009;284:23995–24005.

58. Schindelin, J. et al. Fiji: an open-source platform for biological-image analysis. Nat Methods. 2012;9:676–682.

59. Dobin, A. et al. STAR: ultrafast universal RNA-seq aligner. Bioinformatics. 2013;29:15–21.

60. Liao, Y., Smyth, G. K. & Shi, W. featureCounts: an efficient general purpose program for assigning sequence reads to genomic features. Bioinformatics. 2014;30:923–930.

61. Patro, R., Duggal, G., Love, M. I., Irizarry, R. A. & Kingsford, C. Salmon provides fast and bias-aware quantification of transcript expression. Nat Methods. 2017;14:417–419.

62. Wang, L., Wang, S. & Li, W. RSeQC: quality control of RNA-seq experiments. Bioinformatics. 2012;28:2184–2185.

63. Robinson, M. D., McCarthy, D. J. & Smyth, G. K. edgeR: a Bioconductor package for differential expression analysis of digital gene expression data. Bioinformatics. 2010;26:139–140.

64. Ritchie, M. E. et al. limma powers differential expression analyses for RNA-sequencing and microarray studies. Nucleic Acids Res. 2015;43:e47.

65. Liu, R. et al. Why weight? Modelling sample and observational level variability improves power in RNA-seq analyses. Nucleic Acids Res. 2015;43:e97.

66. Luo, W., Friedman, M. S., Shedden, K., Hankenson, K. D. & Woolf, P. J. GAGE: generally applicable gene set enrichment for pathway analysis. BMC Bioinformatics. 2009;10:161.

67. Zhao, S., Guo, Y., Sheng, Q. & Shyr, Y. Advanced heat map and clustering analysis using heatmap3. Biomed Res Int. 2014;2014:986048.

68. Luo, W. & Brouwer, C. Pathview: an R/Bioconductor package for pathway-based data integration and visualization. Bioinformatics. 2013;29:1830–1831.

69. Cerutti, C. & Ridley, A. J. Endothelial cell-cell adhesion and signaling. Exp Cell Res. 2017;358:31–38.

70. Tanaka-Takiguchi, Y., Kinoshita, M. & Takiguchi, K. Septin-mediated uniform bracing of phospholipid membranes. Curr. Biol. 2009;19:140–145.

71. Bertin, A. et al. Phosphatidylinositol-4,5-bisphosphate promotes budding yeast septin filament assembly and organization. J. Mol. Biol. 2010;404:711–731.

72. Xie, H., Surka, M., Howard, J. & Trimble, W. S. Characterization of the mammalian septin H5: distinct patterns of cytoskeletal and membrane association from other septin proteins. Cell Motil. Cytoskeleton. 1999;43:52–62.

73. Efimova, N. & Svitkina, T. M. Branched actin networks push against each other at adherens junctions to maintain cell-cell adhesion. J. Cell Biol. 2018;217:1827–1845.

74. Takai, Y. & Nakanishi, H. Nectin and afadin: novel organizers of intercellular junctions. J. Cell Sci. 2003;116:17–27.

75. Mandai, K. et al. Afadin: A novel actin filament-binding protein with one PDZ domain localized at cadherin-based cell-to-cell adherens junction. J. Cell Biol. 1997;139:517–528.

76. Okumura, N., Kagami, T., Fujii, K., Nakahara, M. & Koizumi, N. Involvement of Nectin-Afadin in the Adherens Junctions of the Corneal Endothelium. Cornea. 2018;37:633–640.

77. Woodfin, A., Voisin, M. B. & Nourshargh, S. PECAM-1: a multi-functional molecule in inflammation and vascular biology. Arterioscler Thromb Vasc Biol. 2007;27:2514–2523.

78. Zhang, J. et al. Phosphatidylinositol polyphosphate binding to the mammalian septin H5 is modulated by GTP. Curr. Biol. 1999;9:1458–1467.

79. Cannon, K. S., Woods, B. L., Crutchley, J. M. & Gladfelter, A. S. An amphipathic helix enables septins to sense micrometer-scale membrane curvature. J. Cell Biol. 2019;218:1128–1137.

80. Beber, A. et al. Membrane reshaping by micrometric curvature sensitive septin filaments. Nat Commun. 2019;10:420.

81. McLaughlin, S., Wang, J., Gambhir, A. & Murray, D. PIP(2) and proteins: interactions, organization, and information flow. Annu Rev Biophys Biomol Struct. 2002;31:151–175.

82. Goldblum, S. E., Hennig, B., Jay, M., Yoneda, K. & McClain, C. J. Tumor necrosis factor alpha-induced pulmonary vascular endothelial injury. Infect. Immun. 1989;57:1218–1226.

83. Chen, C. C., Sun, Y. T., Chen, J. J. & Chiu, K. T. TNF-alpha-induced cyclooxygenase-2 expression in human lung epithelial cells: involvement of the phospholipase C-gamma 2, protein kinase C-alpha, tyrosine kinase, NF-kappa B-inducing kinase, and I-kappa B kinase 1/2 pathway. J. Immunol. 2000;165:2719–2728.

