## Supplemental table 1 for "Septin Roles and Mechanisms in Organization of Endothelial Cell Junctions"

**Supplemental table 1 Genes that show significant expression change by Septin 2 KD**

| Gene Names | Down (Log2(FC)) | Gene Names | Up (Log2(FC)) |
|---|---|---|---|
| CNR1 | -5.85 | LRRC38 | 5.38 |
| COL1A2 | -5.66 | MMP1 | 5.23 |
| HAPLN1 | -5.50 | PTPRR | 4.96 |
| PLCXD3 | -5.32 | ENKUR | 4.89 |
| TWIST2 | -5.18 | BEST3 | 4.77 |
| SEP2 | -4.64 | MAR11 | 4.64 |
| PRR15 | -4.63 | NCKAP5 | 4.62 |
| SPOCD1 | -4.62 | ADGRG4 | 4.59 |
| NUDT11 | -4.37 | SORL1 | 4.45 |
| ESX1 | -4.21 | NOG | 4.32 |
| VCAN | -4.15 | CYGB | 4.27 |
| RCAN3 | -4.06 | SLC7A14 | 4.02 |
| SLCO4A1 | -3.94 | EPN3 | 3.99 |
| CBR3 | -3.87 | CD22 | 3.98 |
| NEIL3 | -3.86 | KCNH3 | 3.90 |
| CLMP | -3.84 | TMEM236 | 3.71 |
| SLC7A2 | -3.83 | MAR4 | 3.64 |
| CENPM | -3.66 | AL683807.1 | 3.62 |
| UHRF1 | -3.53 | CD226 | 3.58 |
| ENOX1 | -3.49 | C3AR1 | 3.56 |
| CAMK4 | -3.46 | HIST1H2BC | 3.56 |
| G0S2 | -3.44 | COL17A1 | 3.56 |
| SAPCD2 | -3.40 | BCAN | 3.52 |
| CRISPLD2 | -3.39 | JPH2 | 3.49 |
| GBP2 | -3.39 | ELFN2 | 3.40 |
| CDKN2C | -3.38 | ADAM12 | 3.38 |
| COL8A2 | -3.38 | NPAS3 | 3.38 |
| CDC20 | -3.33 | COLQ | 3.37 |
| PKMYT1 | -3.33 | HIF1A-AS2 | 3.36 |
| PCDH10 | -3.31 | AL353740.1 | 3.35 |
| CDC25C | -3.30 | HDAC9 | 3.30 |
| CBLN2 | -3.25 | CLSTN2 | 3.17 |
| FSD1 | -3.24 | LINC01186 | 3.15 |
| UBE2C | -3.23 | XYLT1 | 3.15 |
| GTSE1 | -3.23 | PCDH12 | 3.12 |
| EDIL3 | -3.22 | CLDN3 | 3.04 |
| FAM111B | -3.21 | GATA3 | 3.03 |
| GLIPR1 | -3.21 | SMIM10L2A | 3.02 |
| FAM167A | -3.20 | HIST1H2AD | 2.96 |
| HTR1B | -3.18 | DPEP1 | 2.95 |
| MCM10 | -3.16 | CYTL1 | 2.94 |
| CLSPN | -3.16 | TRAC | 2.89 |

| | | | |
|---|---|---|---|
| ALDH1L2 | -3.13 | TFEC | 2.82 |
| NUF2 | -3.12 | NR4A1 | 2.75 |
| PTGS2 | -3.10 | PGK1.00 | 2.69 |
| TK1 | -3.04 | ACOX2 | 2.68 |
| KCTD16 | -3.03 | HIST1H2BD | 2.64 |
| SHCBP1 | -3.03 | TMC8 | 2.57 |
| PIMREG | -3.00 | IL16 | 2.57 |
| CCNB2 | -2.98 | LOXL4 | 2.50 |
| NDRG4 | -2.98 | AL359643.2 | 2.50 |
| BUB1 | -2.97 | CPM | 2.49 |
| SPC25 | -2.95 | HIST1H2AC | 2.46 |
| CLIC3 | -2.95 | CST6 | 2.46 |
| KIF14 | -2.94 | KCNJ15 | 2.44 |
| RAB20 | -2.94 | GNA15 | 2.44 |
| MYBL2 | -2.94 | TSPAN1 | 2.36 |
| SKA1 | -2.93 | ACER2 | 2.35 |
| NDC80 | -2.93 | CCND2 | 2.33 |
| BMPER | -2.92 | TMC7 | 2.32 |
| FAM129A | -2.92 | WDR66 | 2.31 |
| ERCC6L | -2.91 | HIST1H2BG | 2.30 |
| PLK1 | -2.90 | AL121718.1 | 2.28 |
| SUSD5 | -2.90 | LINC01480 | 2.24 |
| ESCO2 | -2.90 | MMP28 | 2.24 |
| CKAP2L | -2.88 | LINC02356 | 2.23 |
| PBK | -2.86 | FLT1 | 2.22 |
| DLGAP5 | -2.86 | DCBLD2 | 2.21 |
| MEST | -2.86 | ANO4 | 2.20 |
| KIF20A | -2.84 | NOXA1 | 2.14 |
| KIF18B | -2.84 | KRT15 | 2.13 |
| CEP55 | -2.82 | LHX6 | 2.10 |
| HMMR | -2.81 | NR5A2 | 2.10 |
| BLM | -2.80 | LINC01836 | 2.10 |
| AC091057.1 | -2.79 | ZNF429 | 2.08 |
| PLA1A | -2.79 | NT5E | 2.08 |
| KIF4A | -2.78 | SFXN1 | 2.08 |
| ORC1 | -2.77 | MAP2 | 2.05 |
| KHK | -2.77 | LYNX1 | 2.03 |
| SH3BP1 | -2.77 | HIST1H3D | 2.02 |
| WDR62 | -2.76 | LNCOG | 2.01 |
| CDC45 | -2.76 | DGKI | 1.97 |
| CIT | -2.76 | MYLK2 | 1.96 |
| BUB1B | -2.76 | GABRP | 1.95 |
| POLE2 | -2.75 | CFAP57 | 1.94 |
| EXO1 | -2.75 | ACVR2B | 1.93 |
| HJURP | -2.75 | AL139280.1 | 1.93 |

| | | | |
|---|---|---|---|
| RRM2 | -2.74 | CEACAM19 | 1.92 |
| RRM2 | -2.74 | SLC6A4 | 1.91 |
| E2F1 | -2.74 | PCSK6 | 1.89 |
| AURKB | -2.74 | TXLNB | 1.89 |
| E2F8 | -2.74 | ITM2A | 1.88 |
| KIF2C | -2.74 | GPIHBP1 | 1.87 |
| NSG1 | -2.73 | TTC30B | 1.85 |
| KIF15 | -2.72 | BCHE | 1.84 |
| NUSAP1 | -2.72 | SLC16A6 | 1.83 |
| ASF1B | -2.70 | FAM107A | 1.83 |
| BIRC5 | -2.70 | MYO7A | 1.81 |
| NCAPH | -2.70 | SPNS3 | 1.79 |
| RAD54L | -2.69 | CLMN | 1.78 |
| MKI67 | -2.69 | SLC26A2 | 1.78 |
| NEK2 | -2.69 | SLC2A9 | 1.77 |
| CENPA | -2.69 | ARHGEF16 | 1.77 |
| GINS2 | -2.67 | SRPX2 | 1.76 |
| HACD4 | -2.66 | ANKRD46 | 1.76 |
| CDCA8 | -2.66 | ADAM9 | 1.76 |
| HEY2 | -2.65 | KLHL6 | 1.75 |
| ASPM | -2.64 | SULT1A1 | 1.75 |
| SPC24 | -2.63 | PLCXD2 | 1.74 |
| CDCA3 | -2.63 | FUCA1 | 1.73 |
| HSD3B7 | -2.63 | LRP4 | 1.72 |
| DRAXIN | -2.62 | ASTN2 | 1.66 |
| MXD3 | -2.62 | LYRM9 | 1.66 |
| DEPDC1B | -2.61 | HIST1H2BK | 1.66 |
| TTK | -2.61 | CETP | 1.65 |
| KIF18A | -2.61 | HSPA4L | 1.64 |
| TICRR | -2.60 | CD274 | 1.64 |
| BRIP1 | -2.60 | SLC39A6 | 1.62 |
| KNL1 | -2.60 | PPME1 | 1.60 |
| TICAM2 | -2.60 | ITM2C | 1.59 |
| STMN3 | -2.59 | CMTM4 | 1.59 |
| NEO1 | -2.58 | TMEM87B | 1.58 |
| PRR11 | -2.58 | PPM1H | 1.58 |
| TCF19 | -2.58 | AC243960.3 | 1.58 |
| RAD51AP1 | -2.58 | SCAMP5 | 1.58 |
| SLC39A8 | -2.53 | PTPRU | 1.58 |
| TROAP | -2.53 | RNASET2 | 1.57 |
| NCAPG | -2.52 | PDE6A | 1.57 |
| ENTPD6 | -2.52 | PTCHD4 | 1.57 |
| PIM1 | -2.51 | PRKCE | 1.57 |
| MICAL1 | -2.51 | WNT3 | 1.56 |
| C20orf27 | -2.49 | TGFA | 1.55 |

| | | | |
|---|---|---|---|
| IMPA2 | -2.48 | FAS | 1.54 |
| PSRC1 | -2.47 | SESN1 | 1.54 |
| IQGAP3 | -2.46 | TLR1 | 1.54 |
| WDR76 | -2.46 | MEGF6 | 1.53 |
| ARHGAP11A | -2.46 | MYCT1 | 1.53 |
| SLFN11 | -2.45 | OSTM1 | 1.53 |
| GPR162 | -2.44 | RAB11FIP2 | 1.52 |
| PCLAF | -2.43 | KRT80 | 1.52 |
| SGO1 | -2.42 | NEDD4L | 1.50 |
| HASPIN | -2.42 | TNFRSF10C | 1.50 |
| DIXDC1 | -2.41 | MAPRE3 | 1.49 |
| HMOX1 | -2.41 | IL1RAP | 1.49 |
| AKR1C1 | -2.41 | HIST1H1C | 1.48 |
| ANLN | -2.40 | GADD45A | 1.48 |
| CENPE | -2.39 | MERTK | 1.48 |
| CDCA5 | -2.39 | VSIR | 1.47 |
| FKBP1B | -2.38 | UGCG | 1.46 |
| PLK4 | -2.38 | VAV3 | 1.46 |
| MPZL2 | -2.37 | EPB41L1 | 1.46 |
| MELK | -2.37 | HIST1H2AG | 1.46 |
| KREMEN1 | -2.37 | PLAU | 1.46 |
| CENPO | -2.37 | ADGRG6 | 1.45 |
| PTTG1 | -2.36 | BACH2 | 1.45 |
| EHBP1L1 | -2.36 | SRGAP3 | 1.43 |
| CDT1 | -2.36 | ST8SIA4 | 1.43 |
| IL7R | -2.34 | SPATA18 | 1.43 |
| PRC1 | -2.34 | PHTF2 | 1.42 |
| TMEM158 | -2.33 | MIR503HG | 1.42 |
| FOXM1 | -2.33 | MYO1F | 1.41 |
| KIF11 | -2.33 | PIEZO2 | 1.41 |
| SPAG5 | -2.33 | POU2F2 | 1.41 |
| KIF23 | -2.33 | PRNP | 1.40 |
| CCNB1 | -2.33 | ARHGAP25 | 1.40 |
| TOP2A | -2.32 | YOD1 | 1.39 |
| LMNB1 | -2.31 | HIST1H4I | 1.38 |
| CIP2A | -2.30 | CDKN1A | 1.38 |
| CENPI | -2.30 | LINC01094 | 1.38 |
| HHIPL1 | -2.29 | PTPRN2 | 1.38 |
| SKA3 | -2.29 | DCP2 | 1.38 |
| FAM20C | -2.28 | RRM2B | 1.38 |
| CCNA2 | -2.27 | CFAP54 | 1.37 |
| ABCA6 | -2.27 | KLHL28 | 1.37 |
| ADAMTS4 | -2.26 | LYSMD1 | 1.37 |
| POLQ | -2.25 | TNFRSF10B | 1.36 |
| KIFC1 | -2.23 | MRC1 | 1.36 |

| | | | |
|---|---|---|---|
| FBLN5 | -2.23 | ERO1A | 1.36 |
| CDCA2 | -2.22 | LNX1 | 1.36 |
| BASP1 | -2.22 | ZNF493 | 1.36 |
| CHEK1 | -2.21 | KBTBD8 | 1.36 |
| CENPW | -2.20 | SLC2A4 | 1.35 |
| MAD2L1 | -2.19 | B3GLCT | 1.33 |
| KIF24 | -2.18 | E2F5 | 1.33 |
| TPX2 | -2.18 | LINC02188 | 1.33 |
| FANCA | -2.18 | BACH1 | 1.32 |
| CSRP2 | -2.18 | PLA2G4C | 1.32 |
| S1PR3 | -2.17 | LIFR | 1.30 |
| S1PR3 | -2.17 | SLC25A45 | 1.30 |
| MCM4 | -2.17 | ALS2CL | 1.29 |
| CDC6 | -2.17 | TRAPPC6A | 1.29 |
| CXCL5 | -2.17 | CTSD | 1.29 |
| MCM2 | -2.14 | FJX1 | 1.29 |
| NCEH1 | -2.14 | VEGFA | 1.29 |
| PTGER4 | -2.12 | C5orf15 | 1.28 |
| RFC3 | -2.12 | TTC30A | 1.28 |
| CAPG | -2.12 | CPEB3 | 1.27 |
| TMCC2 | -2.11 | PTPRS | 1.27 |
| DPYSL3 | -2.11 | FAM214A | 1.27 |
| APOL4 | -2.09 | CBX7 | 1.27 |
| ESPL1 | -2.09 | SGSM1 | 1.27 |
| ASB13 | -2.09 | SLC25A13 | 1.27 |
| SGO2 | -2.08 | PHACTR2 | 1.27 |
| RACGAP1 | -2.07 | CDKN2B | 1.27 |
| LRRCC1 | -2.06 | MAPK6 | 1.26 |
| CASZ1 | -2.06 | INKA2 | 1.26 |
| LIPG | -2.06 | CDH5 | 1.26 |
| H1FX | -2.05 | GPR143 | 1.26 |
| UBE2T | -2.05 | RIC1 | 1.25 |
| CDC7 | -2.05 | AJUBA | 1.25 |
| STAC | -2.04 | THBD | 1.25 |
| CDK1 | -2.03 | PCMTD1 | 1.25 |
| CENPF | -2.02 | TCAF2 | 1.24 |
| NAALADL1 | -2.02 | CYYR1 | 1.24 |
| PASK | -2.00 | LINC01679 | 1.24 |
| NCAPG2 | -2.00 | DENND1B | 1.23 |
| BRI3BP | -2.00 | CLIP4 | 1.23 |
| FAM83D | -2.00 | GLRB | 1.23 |
| SCARA3 | -1.99 | HOXA1 | 1.22 |
| DEPDC1 | -1.99 | BRWD3 | 1.22 |
| RECQL4 | -1.99 | SLCO2B1 | 1.22 |
| EME1 | -1.99 | REPS2 | 1.22 |

| | | | |
|---|---|---|---|
| FMNL1 | -1.98 | B4GALT4 | 1.21 |
| FAM84A | -1.98 | PLK2 | 1.19 |
| SLC20A1 | -1.98 | DENND5B | 1.19 |
| CTDSP1 | -1.98 | ITGA3 | 1.19 |
| PITPNM1 | -1.96 | INPP4B | 1.19 |
| H2AFY | -1.95 | SLC22A18 | 1.18 |
| SALL2 | -1.94 | HELB | 1.17 |
| GPR37 | -1.93 | PBXIP1 | 1.17 |
| C1orf112 | -1.92 | LRP12 | 1.17 |
| PRDX1 | -1.90 | SLC49A4 | 1.16 |
| CDCA7 | -1.90 | LRG1 | 1.16 |
| ETV4 | -1.88 | FRS2 | 1.16 |
| MEX3B | -1.88 | SLC35G2 | 1.16 |
| MCM6 | -1.88 | ZNF626 | 1.16 |
| OST4 | -1.88 | AMER1 | 1.15 |
| CENPU | -1.88 | ARSB | 1.14 |
| H2AFX | -1.87 | OLFML2A | 1.14 |
| FLNA | -1.87 | DKK3.00 | 1.14 |
| PFKFB3 | -1.87 | C1orf56 | 1.14 |
| TGFB1I1 | -1.87 | DICER1 | 1.13 |
| GINS1 | -1.87 | SDSL | 1.13 |
| CLEC11A | -1.87 | PDE4D | 1.13 |
| AKR1B1 | -1.87 | IL11RA | 1.13 |
| MIEN1 | -1.87 | AK9 | 1.13 |
| ORC6 | -1.86 | TCTN1 | 1.12 |
| OXCT1 | -1.85 | TSKU | 1.12 |
| SUSD2 | -1.85 | DDAH1 | 1.12 |
| BRCA2 | -1.85 | CAVIN2 | 1.12 |
| SRF | -1.85 | LRPAP1 | 1.12 |
| ANKLE1 | -1.84 | SMURF2 | 1.12 |
| PYGB | -1.84 | LDLRAP1 | 1.12 |
| GINS4 | -1.83 | RMND5A | 1.11 |
| LSM14B | -1.83 | HMGA2 | 1.11 |
| TRAIP | -1.83 | MPZL3 | 1.11 |
| STIL | -1.83 | PVR | 1.11 |
| SAT1 | -1.82 | TMEM64 | 1.10 |
| RAC3 | -1.81 | PTP4A3 | 1.10 |
| NAGS | -1.81 | LNPK | 1.10 |
| NPTX2 | -1.81 | STX3 | 1.10 |
| PSMC3IP | -1.80 | LY96 | 1.10 |
| AC103746.1 | -1.80 | C8orf58 | 1.10 |
| B4GALT1 | -1.79 | SGPP1 | 1.10 |
| HELLS | -1.79 | TUSC2 | 1.09 |
| FSCN1 | -1.79 | CREBRF | 1.08 |
| AIFM2 | -1.78 | GALNT7 | 1.08 |

| | | | |
|---|---|---|---|
| UNC13B | -1.78 | MAMLD1 | 1.08 |
| KIF20B | -1.78 | SERINC1 | 1.08 |
| CTSH | -1.78 | CEACAM1 | 1.08 |
| ATP6V0E2 | -1.76 | OCIAD2 | 1.07 |
| RFLNA | -1.76 | RTTN | 1.07 |
| RFLNA | -1.76 | RAMP2-AS1 | 1.06 |
| MID1IP1 | -1.76 | NMT2 | 1.06 |
| DDIAS | -1.75 | STN1.00 | 1.06 |
| DTL | -1.75 | MAN2A2 | 1.06 |
| TENT5A | -1.75 | TMEM68 | 1.05 |
| TRIP13 | -1.74 | DDX3Y | 1.05 |
| IGFBP6 | -1.74 | DLC1 | 1.05 |
| ACOT11 | -1.74 | KLHL24 | 1.05 |
| MGAT5 | -1.74 | ATP9A | 1.04 |
| TCAF1 | -1.72 | ENTPD7 | 1.04 |
| PAQR4 | -1.72 | C1D | 1.04 |
| FNDC11 | -1.72 | PRKAB1 | 1.03 |
| MYRF | -1.72 | GPX8 | 1.03 |
| PC | -1.72 | UHMK1 | 1.03 |
| AGAP2 | -1.71 | DPY19L3 | 1.03 |
| SORD | -1.71 | PALD1 | 1.03 |
| LY75 | -1.70 | SLC39A11 | 1.03 |
| SHMT1 | -1.70 | LCOR | 1.02 |
| ZWINT | -1.70 | LMBR1L | 1.02 |
| SEMA3D | -1.69 | FAM91A1 | 1.02 |
| CHAF1A | -1.69 | MAR9 | 1.02 |
| VEGFC | -1.68 | PTPRE | 1.01 |
| MBOAT1 | -1.68 | NAGLU | 1.01 |
| KIF21B | -1.68 | ZNF462 | 1.01 |
| CALHM2 | -1.67 | BRWD1 | 1.01 |
| WDR4 | -1.66 | SLC16A13 | 1.01 |
| PDLIM1 | -1.65 | RHPN1 | 1.01 |
| LGALS1 | -1.65 | NUDT7 | 1.00 |
| NUDT1 | -1.64 | RAMAC | 1.00 |
| UAP1L1 | -1.63 | SNX4 | 0.99 |
| RBL1 | -1.63 | GNS | 0.99 |
| CBX2 | -1.63 | PPARA | 0.99 |
| CCNF | -1.62 | TMCC1 | 0.99 |
| ZNF703 | -1.61 | MAP3K2 | 0.99 |
| AURKA | -1.61 | TIGAR | 0.98 |
| KIF22 | -1.61 | STRADB | 0.97 |
| CLDN7 | -1.61 | TXNRD2 | 0.97 |
| HMGB3 | -1.61 | ITGB5 | 0.97 |
| PTPN9 | -1.60 | RPS6KA3 | 0.97 |
| ZEB2 | -1.60 | PLCG2 | 0.97 |

| | | | |
|---|---|---|---|
| FUT8 | -1.59 | IL17D | 0.97 |
| HEY1 | -1.59 | C1QTNF6 | 0.96 |
| MBD3 | -1.59 | PRKACB | 0.96 |
| CLIP2 | -1.59 | BTBD3 | 0.96 |
| ATAD2 | -1.59 | ZBTB18 | 0.96 |
| KIAA0513 | -1.58 | BMF | 0.96 |
| GPSM2 | -1.58 | BTD | 0.95 |
| BRCA1 | -1.58 | MAP1LC3B | 0.95 |
| BIN1 | -1.58 | WNK4 | 0.95 |
| GINS3 | -1.58 | MGST2 | 0.95 |
| INCENP | -1.58 | LONRF1 | 0.95 |
| C1QTNF1 | -1.58 | SLC2A10 | 0.95 |
| NACC1 | -1.57 | HIST2H2BE | 0.95 |
| MBOAT2 | -1.57 | RHOBTB1 | 0.95 |
| B3GALT6 | -1.55 | CASP4 | 0.95 |
| PFN1 | -1.54 | KHNYN | 0.95 |
| SRPX | -1.54 | GABARAPL1 | 0.94 |
| SUN2 | -1.54 | OSBP2 | 0.94 |
| VPS51 | -1.54 | ARHGAP45 | 0.94 |
| SCUBE2 | -1.54 | PDIA5 | 0.94 |
| WDHD1 | -1.53 | TM7SF3 | 0.94 |
| LMAN2 | -1.53 | SPTBN5 | 0.93 |
| RAD51 | -1.53 | CD164 | 0.93 |
| TACC3 | -1.53 | IQCK | 0.93 |
| SMPD4 | -1.53 | LMO2 | 0.93 |
| NEXN | -1.53 | OSCP1 | 0.93 |
| SIX1 | -1.52 | GGT1 | 0.93 |
| GCH1 | -1.52 | LATS2 | 0.93 |
| ERMP1 | -1.52 | SLC12A4 | 0.92 |
| OSBPL6 | -1.51 | PPP3CB | 0.92 |
| THG1L | -1.51 | FAM84B | 0.92 |
| DLGAP1 | -1.51 | MBNL2 | 0.92 |
| MTCH2 | -1.50 | TMEM254 | 0.92 |
| SPHK1 | -1.50 | MRS2 | 0.92 |
| MRPS27 | -1.50 | SH3D19 | 0.92 |
| C4orf46 | -1.50 | NDUFA8 | 0.91 |
| PPIH | -1.49 | DOCK7 | 0.91 |
| SNCA | -1.49 | TUFT1 | 0.91 |
| FAM171A1 | -1.49 | YBX3 | 0.91 |
| CDC42EP4 | -1.49 | LMLN | 0.91 |
| LINC00607 | -1.49 | UBN2 | 0.91 |
| LBR | -1.49 | SYCE1L | 0.91 |
| TNFAIP8 | -1.49 | BTBD19 | 0.91 |
| ETS1 | -1.48 | SLC6A8 | 0.91 |
| IDH2 | -1.48 | NF1 | 0.90 |

| | | | |
|---|---|---|---|
| SNHG19 | -1.47 | CLN8 | 0.90 |
| EZH2 | -1.47 | PAFAH2 | 0.90 |
| BARD1 | -1.46 | TMED5 | 0.90 |
| AC080038.1 | -1.44 | ATP7B | 0.90 |
| RPGR | -1.44 | KRT18 | 0.90 |
| C14orf119 | -1.44 | SACM1L | 0.89 |
| ABL1 | -1.44 | THAP6 | 0.89 |
| UBE2S | -1.43 | TMTC3 | 0.89 |
| RMDN1 | -1.43 | ZBTB43 | 0.89 |
| SLC52A2 | -1.43 | ARSD | 0.89 |
| RNF220 | -1.43 | TESK2 | 0.89 |
| PLCB3 | -1.42 | VWA8 | 0.89 |
| ELOVL6 | -1.42 | SLC25A4 | 0.89 |
| DOLPP1 | -1.41 | CPT2 | 0.89 |
| STEAP3 | -1.41 | GYPC | 0.89 |
| FANCI | -1.41 | PPP1R13L | 0.88 |
| PPM1F | -1.41 | SLC35F5 | 0.88 |
| ECI2 | -1.41 | MSRB2 | 0.88 |
| LMNB2 | -1.40 | STK4 | 0.88 |
| VAMP2 | -1.40 | ASAP2 | 0.87 |
| ZNF467 | -1.39 | TEAD1 | 0.87 |
| TRIM59 | -1.39 | MICA | 0.87 |
| TONSL | -1.39 | HTATIP2 | 0.87 |
| MKRN1 | -1.39 | PAQR7 | 0.87 |
| MYBL1 | -1.39 | SLC20A2 | 0.87 |
| AFAP1 | -1.38 | SC5D | 0.86 |
| EPAS1 | -1.38 | TEK | 0.86 |
| CHTF18 | -1.38 | PUDP | 0.86 |
| C12orf75 | -1.37 | TDRP | 0.85 |
| PARP1 | -1.37 | COA5 | 0.85 |
| ARL4D | -1.36 | ZNF43 | 0.85 |
| ANKRD18B | -1.36 | SESN2 | 0.85 |
| TTL | -1.35 | APPL2 | 0.85 |
| CACNB3 | -1.35 | MCFD2 | 0.85 |
| PPP1CA | -1.35 | OTUD4 | 0.85 |
| NAPRT | -1.35 | DCUN1D1 | 0.85 |
| SYT11 | -1.35 | CCPG1 | 0.85 |
| PLAUR | -1.35 | MIGA1 | 0.84 |
| GLIPR2 | -1.35 | MT-TP | 0.84 |
| DPCD | -1.34 | ADARB1 | 0.84 |
| SYTL4 | -1.34 | ASPH | 0.84 |
| NFIC | -1.34 | TMEM132A | 0.84 |
| POC1B | -1.34 | EXOSC5 | 0.84 |
| HIC1 | -1.34 | ECE2 | 0.84 |
| CTSO | -1.34 | CCNDBP1 | 0.84 |

| | | | |
|---|---|---|---|
| MCM5 | -1.34 | ADGRL4 | 0.84 |
| CD99L2 | -1.34 | GDE1 | 0.84 |
| RNASEH2A | -1.34 | SIL1 | 0.84 |
| RAC2 | -1.34 | HIPK3 | 0.83 |
| RFC4 | -1.33 | NEK3 | 0.83 |
| POLD3 | -1.33 | PDGFA | 0.83 |
| LHFPL2 | -1.33 | CCNJ | 0.83 |
| ATP6V1B2 | -1.33 | IMMP2L | 0.83 |
| RUSC1 | -1.33 | GNB4 | 0.83 |
| ARFGAP2 | -1.32 | CARD8 | 0.82 |
| NCAPH2 | -1.31 | RSPH3 | 0.82 |
| CDCA7L | -1.31 | EFNA1 | 0.82 |
| DSN1 | -1.31 | RAB5A | 0.82 |
| TMEM161A | -1.30 | CNEP1R1 | 0.82 |
| RMI2 | -1.30 | SUMF1 | 0.82 |
| SLC35E2B | -1.30 | LIPA | 0.82 |
| CCDC18 | -1.30 | MFSD8 | 0.82 |
| CALM3 | -1.29 | NT5C2 | 0.81 |
| PCSK7 | -1.29 | PIGP | 0.81 |
| FAM89B | -1.29 | AVL9 | 0.81 |
| SNRPA | -1.29 | RALGAPA1 | 0.81 |
| SCLT1 | -1.29 | SLC35D1 | 0.80 |
| PPP1R14B | -1.29 | MLH3 | 0.80 |
| GMPPB | -1.29 | TMEM63B | 0.80 |
| C17orf53 | -1.28 | CDKN1B | 0.80 |
| SSR1 | -1.28 | CYB5R1 | 0.80 |
| GPR176 | -1.28 | CHCHD10 | 0.80 |
| NPM3 | -1.28 | XPC | 0.80 |
| STK38 | -1.28 | ORAI3 | 0.79 |
| MECOM | -1.28 | FKBP4 | 0.79 |
| DYNLL2 | -1.28 | KRT8 | 0.79 |
| IL17RA | -1.28 | MFAP3 | 0.79 |
| DNA2 | -1.27 | ABHD4 | 0.79 |
| CDC25B | -1.27 | PEX5 | 0.79 |
| THOC6 | -1.27 | KLC2 | 0.78 |
| PFAS | -1.27 | ENTPD4 | 0.78 |
| BCL3 | -1.27 | B9D1 | 0.78 |
| PIF1 | -1.27 | CCDC28A | 0.78 |
| NIPAL3 | -1.27 | CCNT2 | 0.78 |
| KLF13 | -1.27 | M6PR | 0.78 |
| STK17B | -1.27 | TSC22D2 | 0.78 |
| ANXA6 | -1.26 | ATAD1 | 0.78 |
| ADCY7 | -1.26 | KDSR | 0.78 |
| MTHFD1 | -1.26 | ATPAF1 | 0.78 |
| NBEAL2 | -1.26 | ABCC1 | 0.77 |

| | | | |
|---|---|---|---|
| TIMM44 | -1.26 | RNF170 | 0.77 |
| MAP3K12 | -1.25 | NDFIP2 | 0.77 |
| SGSH | -1.25 | RB1CC1 | 0.77 |
| TP53I13 | -1.25 | UTP25 | 0.76 |
| DBF4B | -1.25 | ZCCHC14 | 0.76 |
| MARVELD2 | -1.25 | CRBN | 0.76 |
| RAB27A | -1.24 | RGL2 | 0.76 |
| TMCO3 | -1.24 | NRROS | 0.76 |
| C19orf71 | -1.24 | PLCL2 | 0.76 |
| SIPA1L1 | -1.24 | GNPTG | 0.76 |
| APOOL | -1.23 | KDM3A | 0.76 |
| DNMT1 | -1.23 | CRADD | 0.76 |
| MARVELD1 | -1.22 | XKR8 | 0.76 |
| A1BG | -1.22 | KCTD11 | 0.75 |
| USP1 | -1.22 | FKTN | 0.75 |
| FEN1 | -1.22 | PHLPP2 | 0.75 |
| RAB3B | -1.22 | SMIM20 | 0.75 |
| SERPINH1 | -1.22 | CYBRD1 | 0.75 |
| CMTM3 | -1.22 | PCNX4 | 0.75 |
| NCBP3 | -1.22 | KCTD20 | 0.75 |
| MMP19 | -1.22 | VTA1 | 0.75 |
| PMM1 | -1.22 | DNAJC16 | 0.75 |
| PSKH1 | -1.22 | NBR1 | 0.74 |
| EPS8L1 | -1.22 | MINPP1 | 0.74 |
| CLCF1 | -1.22 | GLO1 | 0.74 |
| DNAJC9 | -1.22 | FBXL20 | 0.74 |
| RELT | -1.22 | DNAJB9 | 0.74 |
| AP3B1 | -1.22 | KLC4 | 0.74 |
| EFNB2 | -1.21 | DAGLA | 0.74 |
| TMEM106C | -1.21 | ESAM | 0.74 |
| INSIG1 | -1.21 | KIF3B | 0.74 |
| MAP3K5 | -1.21 | TPRG1L | 0.73 |
| GALNT15 | -1.21 | VPS4B | 0.73 |
| RAPH1 | -1.21 | NCSTN | 0.73 |
| WDR34 | -1.20 | BSDC1 | 0.72 |
| PDCD1LG2 | -1.20 | EFR3A | 0.72 |
| POTEF | -1.20 | SYS1 | 0.72 |
| TLNRD1 | -1.19 | TMEM59 | 0.72 |
| ILF3-DT | -1.19 | CHST12 | 0.72 |
| MADD | -1.19 | MAN1A2 | 0.72 |
| TRMT112 | -1.19 | GOLGA2 | 0.71 |
| NAMPTP1 | -1.19 | MOCS3 | 0.71 |
| ZNF512 | -1.19 | SLC9A1 | 0.71 |
| RCC1L | -1.19 | TRMT6 | 0.71 |
| CHN1 | -1.19 | ACVR1B | 0.71 |

| | | | |
|---|---|---|---|
| UBXN2B | -1.18 | ATP6V1G1 | 0.71 |
| LRP8 | -1.18 | TMX2 | 0.71 |
| CENPJ | -1.18 | PTPN4 | 0.71 |
| MCUB | -1.18 | NECAP1 | 0.70 |
| PNPO | -1.18 | YTHDF3 | 0.70 |
| MICAL2 | -1.18 | TMEM106B | 0.70 |
| MICAL2 | -1.18 | SYNJ1 | 0.70 |
| MAP3K6 | -1.18 | SLC30A6 | 0.70 |
| FAM53B | -1.16 | LRRC75A | 0.70 |
| ECH1 | -1.16 | RPS6KC1 | 0.70 |
| SFRP1 | -1.16 | LIMD1 | 0.69 |
| ARPC4 | -1.16 | SERTAD1 | 0.69 |
| FAAP100 | -1.16 | POLD4 | 0.69 |
| ARHGDIA | -1.16 | WLS | 0.69 |
| PRKD1 | -1.16 | CYTH3 | 0.69 |
| GIMAP7 | -1.16 | PRUNE2 | 0.69 |
| SBF1 | -1.15 | JMY | 0.68 |
| NCAPD3 | -1.15 | TRAM2 | 0.68 |
| CALU | -1.15 | RUFY3 | 0.68 |
| ZNF785 | -1.15 | ZNF800 | 0.68 |
| NME3 | -1.15 | SMIM13 | 0.68 |
| ADGRA2 | -1.15 | ATP7A | 0.68 |
| RNF24 | -1.15 | ERGIC2 | 0.68 |
| PIGC | -1.15 | ERLIN2 | 0.67 |
| SLC25A22 | -1.15 | NHLRC2 | 0.67 |
| TYMP | -1.15 | MIA3 | 0.67 |
| POLD1 | -1.14 | DYRK1B | 0.66 |
| FAM171A2 | -1.14 | NMRK1 | 0.66 |
| VARS | -1.14 | GAA | 0.66 |
| NAMPT | -1.14 | CBWD2 | 0.66 |
| PTGES2 | -1.14 | ZNF268 | 0.66 |
| DAXX | -1.14 | IFNAR1 | 0.65 |
| DHRS4 | -1.13 | PELO | 0.65 |
| CBX6 | -1.13 | C1GALT1 | 0.65 |
| DNAJC5 | -1.13 | STMP1 | 0.65 |
| NUDT3 | -1.12 | DNAL1 | 0.65 |
| MLXIP | -1.12 | ICAM2 | 0.65 |
| NME4 | -1.12 | ANKIB1 | 0.65 |
| LPCAT1 | -1.12 | SUN1 | 0.64 |
| SUPT6H | -1.11 | MRRF | 0.64 |
| SESTD1 | -1.11 | DNAJB12 | 0.64 |
| KDM1A | -1.11 | TCAIM | 0.64 |
| LRFN3 | -1.11 | PIAS2 | 0.64 |
| SFT2D1 | -1.11 | EIF4G3 | 0.64 |
| CRNDE | -1.11 | LEMD2 | 0.64 |

| | | | |
|---|---|---|---|
| HPSE | -1.11 | UNC50 | 0.64 |
| WDR54 | -1.10 | MBTPS2 | 0.64 |
| MAP3K13 | -1.10 | SLFN12 | 0.64 |
| CCR10 | -1.10 | RTCA | 0.63 |
| REPIN1 | -1.10 | RBM18 | 0.63 |
| ENO2 | -1.10 | SETD7 | 0.63 |
| NCKIPSD | -1.10 | STX17 | 0.63 |
| INTS9 | -1.10 | CDYL | 0.63 |
| DHRS4L2 | -1.10 | RHOBTB3 | 0.63 |
| PYCR3 | -1.10 | PRMT3 | 0.63 |
| PLP2 | -1.10 | ANXA3 | 0.63 |
| MVK | -1.09 | CMPK1 | 0.63 |
| SUFU | -1.09 | ADAM17 | 0.62 |
| ARL6IP1 | -1.09 | ERCC1 | 0.62 |
| ODC1 | -1.09 | SERINC3 | 0.62 |
| DAZAP1 | -1.08 | RIT1 | 0.62 |
| CHST7 | -1.08 | PLXNB1 | 0.62 |
| C15orf61 | -1.08 | CPPED1 | 0.62 |
| TTLL12 | -1.08 | SLC7A6 | 0.62 |
| STARD10 | -1.08 | ERCC5 | 0.62 |
| LRBA | -1.08 | ERCC5 | 0.62 |
| CCDC71L | -1.07 | UBE3B | 0.62 |
| ZNF48 | -1.07 | PPM1A | 0.61 |
| GLUL | -1.07 | C6orf120 | 0.61 |
| RCAN1 | -1.07 | DDX1 | 0.61 |
| TAGLN2 | -1.07 | ERCC4 | 0.60 |
| ARPIN | -1.07 | TKFC | 0.60 |
| SLC37A4 | -1.07 | FAM214B | 0.60 |
| NAXD | -1.07 | CHMP5 | 0.60 |
| MYDGF | -1.07 | PCAT19 | 0.60 |
| MFAP2 | -1.07 | TMEM9 | 0.59 |
| FANCG | -1.07 | PPP1R15B | 0.59 |
| PRPS2 | -1.07 | RNF6 | 0.59 |
| NRM | -1.06 | RHOQ | 0.59 |
| CTSC | -1.06 | SHTN1 | 0.59 |
| MEX3A | -1.06 | BAK1 | 0.59 |
| AP001505.1 | -1.06 | NDUFB5 | 0.59 |
| TNPO1 | -1.06 | CNOT6 | 0.59 |
| SLC39A3 | -1.06 | LDB1 | 0.59 |
| FSTL3 | -1.06 | MOSPD2 | 0.59 |
| BNC1 | -1.06 | UBE2H | 0.59 |
| GPR161 | -1.06 | ATP6V1D | 0.59 |
| ZNF746 | -1.06 | LZIC | 0.58 |
| AP2S1 | -1.06 | EMC7 | 0.58 |
| RANBP1 | -1.06 | HSD17B12 | 0.58 |

| | | | |
|---|---|---|---|
| PITPNA | -1.06 | SNAPC5 | 0.58 |
| FZD1 | -1.06 | CHMP1B | 0.58 |
| ACOT8 | -1.05 | CRY2 | 0.58 |
| C11orf95 | -1.05 | SPR | 0.58 |
| VMA21 | -1.05 | MICB | 0.58 |
| FECH | -1.05 | IGF2BP3 | 0.57 |
| PTMS | -1.05 | APH1B | 0.57 |
| UBR7 | -1.05 | NEK6 | 0.57 |
| AC092171.2 | -1.04 | ZMAT3 | 0.57 |
| AP1M1 | -1.04 | SWI5 | 0.57 |
| HNRNPD | -1.04 | NAPG | 0.57 |
| THOC3 | -1.04 | GGA1 | 0.57 |
| WASF3 | -1.04 | AHR | 0.57 |
| SRSF2 | -1.04 | GID4 | 0.56 |
| JKAMP | -1.04 | NR1D2 | 0.56 |
| EXTL3 | -1.03 | PDZD8 | 0.56 |
| TOMM34 | -1.03 | PIK3R3 | 0.56 |
| DGCR2 | -1.03 | CTR9 | 0.56 |
| ZNF205 | -1.03 | GMFG | 0.56 |
| NYAP1 | -1.03 | TTC19 | 0.56 |
| TRAPPC9 | -1.03 | GTF3C3 | 0.56 |
| SAMD10 | -1.02 | GOLGA4 | 0.56 |
| LINC00987 | -1.02 | SENP2 | 0.55 |
| TTLL11 | -1.02 | MGME1 | 0.55 |
| AGPAT2 | -1.02 | FXR1 | 0.55 |
| EIF2AK1 | -1.02 | OCIAD1 | 0.55 |
| ALDH16A1 | -1.01 | AC010618.1 | 0.54 |
| CNOT9 | -1.01 | CLN5 | 0.54 |
| EIF4EBP1 | -1.01 | SIRT2 | 0.54 |
| NSD2 | -1.01 | POFUT1 | 0.54 |
| RBFOX2 | -1.01 | SRA1 | 0.54 |
| ERI1 | -1.01 | TMEM167B | 0.54 |
| CEP89 | -1.01 | SPPL2A | 0.54 |
| SLC45A3 | -1.01 | SCD5 | 0.53 |
| ASL | -1.00 | GIPC1 | 0.53 |
| SEMA6B | -1.00 | ANAPC13 | 0.53 |
| HAUS2 | -1.00 | FHL3 | 0.53 |
| PDE4DIP | -1.00 | CERS2 | 0.53 |
| TACC1 | -1.00 | HDAC1 | 0.53 |
| ALDH9A1 | -1.00 | TMEM9B | 0.53 |
| LRFN4 | -0.99 | RARS2 | 0.52 |
| RTL8C | -0.99 | RDH14 | 0.52 |
| CORO1C | -0.99 | RTL10 | 0.52 |
| MVD | -0.99 | TAX1BP1 | 0.52 |
| FBXL18 | -0.98 | NTPCR | 0.52 |

| | | | |
|---|---|---|---|
| SMC4 | -0.98 | TBCK | 0.52 |
| C16orf45 | -0.98 | MTMR3 | 0.52 |
| TPRN | -0.98 | HEG1 | 0.50 |
| EGLN2 | -0.98 | MAP2K4 | 0.50 |
| PSEN2 | -0.97 | MTMR9 | 0.50 |
| SAP30BP | -0.97 | PDHX | 0.50 |
| AP3S2 | -0.97 | CAPN7 | 0.50 |
| UBAC1 | -0.97 | TRPC4AP | 0.49 |
| PRUNE1 | -0.97 | CSNK1G3 | 0.48 |
| FARSA | -0.96 | TM9SF4 | 0.48 |
| TXNDC15 | -0.96 | IARS2 | 0.48 |
| NDUFA9 | -0.96 | SNX11 | 0.48 |
| AC080112.1 | -0.96 | HBP1 | 0.47 |
| BLOC1S4 | -0.96 | PCNP | 0.46 |
| SLC12A7 | -0.96 | ANKRD40 | 0.46 |
| TSPAN14 | -0.96 | TCEA1 | 0.45 |
| KCTD12 | -0.96 | SS18 | 0.45 |
| ARHGAP22 | -0.95 | CHFR | 0.44 |
| FOXK1 | -0.95 | | |
| WTAP | -0.95 | | |
| EXOSC6 | -0.95 | | |
| PAXX | -0.95 | | |
| INTS13 | -0.95 | | |
| CNNM4 | -0.95 | | |
| TMEM104 | -0.95 | | |
| GOLM1 | -0.95 | | |
| NDST1 | -0.95 | | |
| DFFA | -0.94 | | |
| CDK2 | -0.94 | | |
| MIS18BP1 | -0.94 | | |
| RNF126 | -0.94 | | |
| MACROD1 | -0.94 | | |
| ADIPOR1 | -0.94 | | |
| LLGL1 | -0.94 | | |
| FAM126A | -0.94 | | |
| ADH5 | -0.93 | | |
| KIF3C | -0.93 | | |
| SURF4 | -0.93 | | |
| RHOT2 | -0.93 | | |
| G6PD | -0.93 | | |
| VAT1 | -0.93 | | |
| TBC1D16 | -0.93 | | |
| UBFD1 | -0.93 | | |
| FOXP1 | -0.93 | | |
| BOP1.00 | -0.92 | | |

| | |
|---|---|
| NIPA2 | -0.92 |
| FAM234A | -0.92 |
| STARD7 | -0.92 |
| NCOA7 | -0.92 |
| RAB8B | -0.92 |
| PARVA | -0.92 |
| WRAP53 | -0.92 |
| SH3PXD2B | -0.91 |
| TRABD | -0.91 |
| RBM10 | -0.91 |
| ICMT | -0.91 |
| MEX3D | -0.91 |
| TOR3A | -0.91 |
| H2AFV | -0.91 |
| MRPS12 | -0.90 |
| TCEAL3 | -0.90 |
| UNKL | -0.90 |
| CIAO3 | -0.90 |
| ARF5 | -0.89 |
| ATP6V1A | -0.89 |
| CCDC88A | -0.89 |
| PLPP3 | -0.89 |
| BCAT2 | -0.89 |
| SUMO3 | -0.88 |
| DTYMK | -0.88 |
| TRIM65 | -0.88 |
| ANKRD9 | -0.88 |
| STOML1 | -0.88 |
| INPP5F | -0.88 |
| NR2F2 | -0.88 |
| MAP3K1 | -0.88 |
| ZFP36L1 | -0.88 |
| NUMBL | -0.87 |
| EML3 | -0.87 |
| MGST1 | -0.87 |
| SMAD3 | -0.87 |
| BAZ1A | -0.87 |
| MSRB1 | -0.87 |
| TPM4 | -0.87 |
| FAR2 | -0.86 |
| POLR3H | -0.86 |
| TMCC3 | -0.86 |
| FAM120A | -0.86 |
| TTYH3 | -0.86 |
| B3GNT2 | -0.86 |

| | |
|---|---|
| PGAM1 | -0.85 |
| NARF | -0.85 |
| RBMX2 | -0.85 |
| SPATA2L | -0.85 |
| CNTNAP1 | -0.85 |
| CXXC1 | -0.85 |
| SEC62 | -0.85 |
| ARL10.00 | -0.85 |
| YME1L1 | -0.85 |
| KDELC2 | -0.84 |
| PPP3CC | -0.84 |
| MCM3 | -0.84 |
| FPGS | -0.84 |
| CNTLN | -0.84 |
| ZBED1 | -0.84 |
| KLHL42 | -0.84 |
| RASIP1 | -0.83 |
| DCAF7 | -0.83 |
| ACOT7 | -0.83 |
| TBC1D14 | -0.83 |
| CEP78 | -0.83 |
| ZDHHC7 | -0.83 |
| PIAS4 | -0.83 |
| NABP2 | -0.83 |
| SPECC1L | -0.83 |
| IER5 | -0.83 |
| SNHG9 | -0.83 |
| AP1B1 | -0.82 |
| LARP4B | -0.82 |
| MIEF1 | -0.82 |
| HDHD5 | -0.82 |
| ACLY | -0.82 |
| AL162171.1 | -0.82 |
| EVA1C | -0.82 |
| SMAP2 | -0.81 |
| CHTF8 | -0.81 |
| CHTF8 | -0.81 |
| PRXL2B | -0.81 |
| UGDH | -0.81 |
| GAB2 | -0.81 |
| PCYT2 | -0.81 |
| SEP9 | -0.81 |
| ANKFY1 | -0.80 |
| AKTIP | -0.80 |
| MAPKAPK2 | -0.80 |

| | |
|---|---|
| PUF60 | -0.80 |
| ALDH1B1 | -0.80 |
| UBTD2 | -0.80 |
| ALMS1 | -0.80 |
| TMEM260 | -0.80 |
| ASF1A | -0.80 |
| BCL6 | -0.80 |
| ZNF512B | -0.79 |
| TAL1 | -0.79 |
| ZYX | -0.79 |
| DNAJC10 | -0.79 |
| SLC2A4RG | -0.79 |
| DOCK9 | -0.79 |
| SHQ1 | -0.79 |
| SSRP1 | -0.79 |
| VMP1 | -0.79 |
| PKD1 | -0.79 |
| KLHL5 | -0.79 |
| NCDN | -0.78 |
| SGF29 | -0.78 |
| PQLC1 | -0.78 |
| ZZEF1 | -0.78 |
| STK25 | -0.78 |
| GFOD1 | -0.78 |
| KIAA0586 | -0.78 |
| PXN | -0.77 |
| ANP32E | -0.77 |
| LRRC8A | -0.77 |
| GTF2IRD1 | -0.77 |
| FAM129B | -0.77 |
| CTU2 | -0.77 |
| POP7 | -0.77 |
| AKAP1 | -0.77 |
| ARMC6 | -0.76 |
| AAAS | -0.76 |
| RILPL2 | -0.76 |
| BLOC1S3 | -0.76 |
| TNK2 | -0.76 |
| ZNF775 | -0.76 |
| USP22 | -0.76 |
| ACTR2 | -0.76 |
| RCN3 | -0.75 |
| POLDIP2 | -0.75 |
| DBR1 | -0.75 |
| USP19 | -0.75 |

| | |
|---|---|
| RTL8B | -0.75 |
| ABCC10 | -0.75 |
| JMJD8 | -0.75 |
| ADAMTSL1 | -0.74 |
| TCTA | -0.74 |
| PRR14 | -0.74 |
| LSM14A | -0.74 |
| BCS1L | -0.74 |
| SGTA | -0.74 |
| RTL8A | -0.74 |
| ZBTB45 | -0.74 |
| MAPK14 | -0.74 |
| TRAFD1 | -0.74 |
| TIGD5 | -0.74 |
| ZBTB47 | -0.74 |
| MGAT4B | -0.73 |
| GRB10 | -0.73 |
| EIF2B5 | -0.73 |
| LHFPL6 | -0.73 |
| EIF4H | -0.73 |
| CCM2 | -0.73 |
| SLC22A23 | -0.73 |
| ORAI2 | -0.73 |
| NAV2 | -0.73 |
| PIGS | -0.73 |
| PRR12 | -0.72 |
| NGRN | -0.72 |
| ZNF330 | -0.72 |
| SLC16A1 | -0.72 |
| SH3TC1 | -0.72 |
| CAVIN1 | -0.72 |
| CSNK1G2 | -0.72 |
| STMN1 | -0.72 |
| C8orf33 | -0.72 |
| SLC39A13 | -0.72 |
| WIPI2 | -0.72 |
| KLF16 | -0.71 |
| ST3GAL2 | -0.71 |
| SNX8 | -0.71 |
| TOR4A | -0.71 |
| PJA1 | -0.71 |
| DHX37 | -0.71 |
| UBE2I | -0.71 |
| DTD1 | -0.71 |
| SQOR | -0.71 |

| | |
|---|---|
| FADD | -0.71 |
| MARK3 | -0.71 |
| DPH7 | -0.70 |
| TSPYL1 | -0.70 |
| MAP4K5 | -0.70 |
| ALYREF | -0.70 |
| CTDSP2 | -0.70 |
| NDE1 | -0.70 |
| TTC38 | -0.70 |
| VPS37C | -0.70 |
| GRB2 | -0.70 |
| RCOR1 | -0.70 |
| CCNG2 | -0.70 |
| FP565260.2 | -0.70 |
| CELF1 | -0.70 |
| MLST8 | -0.70 |
| C19orf54 | -0.70 |
| MAP4 | -0.70 |
| TULP3 | -0.70 |
| BAZ1B | -0.70 |
| SLC29A1 | -0.70 |
| CARS2 | -0.69 |
| CACNA2D1 | -0.69 |
| NCLN | -0.69 |
| TPGS2 | -0.69 |
| PAPSS1 | -0.69 |
| TCF20 | -0.69 |
| AXIN1 | -0.68 |
| SMC1A | -0.68 |
| IP6K1 | -0.68 |
| MKRN2 | -0.68 |
| ZNF428 | -0.68 |
| FBXO46 | -0.68 |
| MAD2L1BP | -0.68 |
| LRRC47 | -0.68 |
| CNRIP1 | -0.68 |
| GPD2 | -0.68 |
| TOP3A | -0.68 |
| RUBCN | -0.68 |
| DCAKD | -0.67 |
| POLRMT | -0.67 |
| FZR1 | -0.67 |
| SART1 | -0.67 |
| MED16 | -0.67 |
| CSNK1G1 | -0.67 |

| | |
|---|---|
| WDTC1 | -0.67 |
| TCF7L1 | -0.66 |
| CANT1 | -0.66 |
| CPT1A | -0.66 |
| CLUH | -0.66 |
| GRK6 | -0.66 |
| DIP2B | -0.66 |
| SLC10A3 | -0.66 |
| PKM | -0.65 |
| TOPBP1 | -0.65 |
| FBXO31 | -0.65 |
| NES | -0.65 |
| BAHCC1 | -0.65 |
| MAP4K4 | -0.64 |
| CAD | -0.64 |
| TIMM50 | -0.64 |
| ARHGAP35 | -0.64 |
| TUBB6 | -0.64 |
| DDAH2 | -0.64 |
| PTTG1IP | -0.63 |
| SSU72 | -0.63 |
| DPM3 | -0.63 |
| ZNF74 | -0.63 |
| PISD | -0.63 |
| SLC25A39 | -0.63 |
| MAFG | -0.63 |
| TRIM3 | -0.63 |
| HDAC6 | -0.63 |
| PRMT2 | -0.62 |
| CHST2 | -0.62 |
| ATN1 | -0.62 |
| RALY | -0.62 |
| CHID1 | -0.62 |
| NPLOC4 | -0.62 |
| SLC25A11 | -0.62 |
| ARL4A | -0.62 |
| BRD4 | -0.62 |
| NLGN2 | -0.62 |
| SPOUT1 | -0.62 |
| CMAS | -0.61 |
| GRK2 | -0.61 |
| SLC35A4 | -0.61 |
| FBXO7 | -0.61 |
| SIPA1 | -0.61 |
| MTDH | -0.61 |

| | |
|---|---|
| NELFB | -0.60 |
| CCDC124 | -0.60 |
| CREB3L2 | -0.60 |
| PPP4C | -0.60 |
| RSPRY1 | -0.60 |
| POM121 | -0.60 |
| CHERP | -0.60 |
| CRKL | -0.60 |
| MRPL28 | -0.59 |
| IDH1 | -0.59 |
| GIMAP5 | -0.59 |
| STRN4 | -0.59 |
| PHB | -0.59 |
| PIGQ | -0.59 |
| MECP2 | -0.59 |
| ZMIZ2 | -0.59 |
| SF3A3 | -0.59 |
| SERPINB9 | -0.59 |
| RNF19B | -0.58 |
| RPA1 | -0.58 |
| DCTPP1 | -0.58 |
| DBNL | -0.58 |
| WDR45B | -0.58 |
| GTPBP1 | -0.57 |
| LMF2 | -0.57 |
| CYBA | -0.57 |
| COPS7B | -0.57 |
| DIAPH1 | -0.57 |
| BRK1 | -0.57 |
| UPF1 | -0.57 |
| CTBP1 | -0.57 |
| RHOG | -0.56 |
| SH3RF3 | -0.56 |
| NDUFV1 | -0.56 |
| YWHAQ | -0.56 |
| UBL4A | -0.56 |
| SHISA5 | -0.56 |
| TCF3 | -0.56 |
| ANP32A | -0.56 |
| ZBTB1 | -0.55 |
| OGFOD3 | -0.55 |
| SAFB2 | -0.55 |
| GIMAP8 | -0.55 |
| WDR6 | -0.55 |
| BTBD2 | -0.55 |

| | |
|---|---|
| IVNS1ABP | -0.55 |
| MUS81 | -0.55 |
| B3GALNT2 | -0.54 |
| LRCH3 | -0.54 |
| MCRIP1 | -0.54 |
| GRWD1 | -0.54 |
| RNH1 | -0.54 |
| SNAP47 | -0.54 |
| ARID1B | -0.54 |
| ARMC5 | -0.53 |
| RNF34 | -0.53 |
| ABCB8 | -0.53 |
| ACO2 | -0.53 |
| PSMD5 | -0.52 |
| EXD2 | -0.52 |
| MON1B | -0.52 |
| ZFYVE1 | -0.51 |
| NNT | -0.51 |
| YY1AP1 | -0.51 |
| SEC22C | -0.50 |
| DLST | -0.49 |
| ASXL1 | -0.49 |
| RAP2B | -0.48 |
| SNX1 | -0.48 |
| RSU1 | -0.48 |
| NUBP2 | -0.48 |
| ASB6 | -0.47 |
| RER1 | -0.47 |
| LETM1 | -0.47 |
| USF1 | -0.46 |
| CHMP7 | -0.45 |
| PPP2R1A | -0.45 |
| ATXN7L3 | -0.45 |
| PCNX3 | -0.44 |
| RAB35 | -0.44 |
