## Supplemental table 2 for "Septin Roles and Mechanisms in Organization of Endothelial Cell Junctions"

**Supplement table 2. Genes that show significant expression change by TNF-a**

| Gene Names | Down (log2(FC)) | Gene Names | Up (log2(FC)) |
|---|---|---|---|
| CCL1 | -3.18 | U2AF1 | 1.34 |
| FOXF1 | -1.72 | BEST1 | 1.30 |
| WHRN | -1.39 | CXCL11 | 1.08 |
| FAAH | -1.05 | AC007383.2 | 0.97 |
| SPATA13 | -1.01 | FBXO41 | 0.87 |
| ZNF443 | -0.94 | LINC00513 | 0.83 |
| TYW1B | -0.80 | TRAF1 | 0.81 |
| ZNF433 | -0.78 | CEACAM16 | 0.74 |
| ZNF230 | -0.73 | Sept1 | 0.68 |
| IQCA1 | -0.72 | GAL3ST4 | 0.63 |
| ZNF684 | -0.69 | PCDHB8 | 0.62 |
| AMOT | -0.64 | LDLRAD2 | 0.61 |
| ZNF624 | -0.54 | MIR126 | 0.58 |
| AC103746.1 | -0.52 | AC006001.3 | 0.56 |
| SCUBE2 | -0.46 | SPTBN5 | 0.55 |
| TBC1D19 | -0.45 | MIR222HG | 0.52 |
| WDSUB1 | -0.45 | NEAT1 | 0.52 |
| CBR4 | -0.37 | DPF3 | 0.52 |
| SOCS5 | -0.34 | KIF16B | 0.47 |
| RHOU | -0.33 | GOLGA8A | 0.46 |
| AC026785.2 | -0.32 | GABRE | 0.43 |
| BBS9 | -0.27 | SMN2 | 0.40 |
| LMBRD1 | -0.25 | LINC00346 | 0.40 |
| TMEM87A | -0.24 | NPIPB4 | 0.39 |
| ELF1 | -0.23 | KCTD13 | 0.39 |
| ASB8 | -0.23 | RABGEF1 | 0.36 |
| VPS37A | -0.21 | COL27A1 | 0.33 |
| | | BTBD19 | 0.33 |
| | | CBWD5 | 0.31 |
| | | MYEF2 | 0.30 |
| | | EP400P1 | 0.28 |
| | | MIR22HG | 0.26 |
| | | HIP1R | 0.26 |
| | | DNAJC11 | 0.24 |
| | | PNN | 0.23 |
| | | CFLAR | 0.20 |
