## Supplemental table 3 for "Septin Roles and Mechanisms in Organization of Endothelial Cell Junctions"

**Supplemental table 3 Changes of Intercellular adhesion molecules by septin 2 KD**

| Gene name | log2 (FC) | negative log10 (adj.P.Value) |
|---|---|---|
| CDH5 ** | 1.26 | 1.8 |
| CTNNB1 | 0.91 | 1.12 |
| CTNNA1 | 0.13 | 0.23 |
| CTNND1 | 0.02 | 0.02 |
| NECTIN1 | 0.43 | 0.42 |
| NECTIN2 ** | 0.27 | 0.59 |
| NECTIN3 | -0.08 | 0.07 |
| AFDN ** | 0.00 | 0 |
| PECAM1 ** | 0.68 | 0.91 |
| TJP1 ** | 0.09 | 0.07 |
| TJP2 | 0.24 | 0.34 |
| OCLN | -0.28 | 0.12 |
| CLDN1 | -0.92 | 0.7 |
| CLDN3 | 3.04 | 1.51 |
| CLDN4 | 1.10 | 0.72 |
| CLDN5 | 0.54 | 0.21 |
| CLDN7 | -1.61 | 1.37 |
| CLDN10 | -4.66 | 0.89 |
| CLDN11 | -0.29 | 0.12 |
| CLDN12 | 0.38 | 0.51 |
| CLDN14 | 0.96 | 0.87 |
| CLDN15 | 0.14 | 0.1 |
| JAM3 | 0.63 | 0.78 |
| CD99 | -0.35 | 0.73 |
| ACTB | -0.76 | 0.88 |
| ACTA1 | -0.79 | 0.57 |
| ACTA2 | 1.03 | 1.18 |
| ACTG1 | -0.20 | 0.18 |
