## Supplemental table 4 for "Septin Roles and Mechanisms in Organization of Endothelial Cell Junctions"

**Supplemental table 4 Changes of intercellular adhesion molecules by TNFα treatment**

| Gene name | log2 (FC) | negative log10 (P.Value) |
|---|---|---|
| CDH5 ** | 0.09 | 0.21 |
| CTNNA1 | 0.04 | 0.15 |
| CTNNB1 | -0.02 | 0.03 |
| CTNND1 | 0.03 | 0.09 |
| NECTIN1 | -0.08 | 0.15 |
| NECTIN2 ** | 0.01 | 0.03 |
| NECTIN3 | -0.06 | 0.15 |
| AFDN ** | -0.03 | 0.1 |
| PECAM1 ** | -0.03 | 0.07 |
| TJP1 ** | -0.05 | 0.12 |
| TJP2 | 0.04 | 0.12 |
| OCLN | 0.21 | 0.26 |
| CLDN11 | 0.15 | 0.17 |
| CLDN15 | 0.06 | 0.09 |
| CLDN10 | -0.23 | 0.1 |
| CLDN12 | -0.12 | 0.3 |
| CLDN14 | 0.31 | 0.52 |
| CLDN1 | 0.38 | 0.44 |
| CLDN3 | 0.20 | 0.16 |
| CLDN7 | 0.08 | 0.12 |
| CLDN5 | -0.25 | 0.23 |
| CLDN4 | -0.11 | 0.1 |
| CD99 | -0.07 | 0.26 |
| JAM3 | -0.13 | 0.3 |
| ACTB | 0.14 | 0.28 |
| ACTA2 | -0.02 | 0.03 |
| ACTA1 | 0.19 | 0.26 |
| ACTG1 | 0.10 | 0.25 |
